# Identifying Gene Expression Programs of Cell-type Identity and Cellular Activity with Single-Cell RNA-Seq

**DOI:** 10.1101/310599

**Authors:** Dylan Kotliar, Adrian Veres, M. Aurel Nagy, Shervin Tabrizi, Eran Hodis, Douglas A. Melton, Pardis C. Sabeti

## Abstract

Identifying gene expression programs underlying both cell-type identity and cellular activities (e.g. life-cycle processes, responses to environmental cues) is crucial for understanding the organization of cells and tissues. Although single-cell RNA-Seq (scRNA-Seq) can quantify transcripts in individual cells, each cell’s expression profile may be a mixture of both types of programs, making them difficult to disentangle. Here we illustrate and enhance the use of matrix factorization as a solution to this problem. We show with simulations that a method that we call consensus non-negative matrix factorization (cNMF) accurately infers identity and activity programs, including the relative contribution of programs in each cell. Applied to published brain organoid and visual cortex scRNA-Seq datasets, cNMF refines the hierarchy of cell types and identifies both expected (e.g. cell cycle and hypoxia) and intriguing novel activity programs. We propose that one of the novel programs may reflect a neurosecretory phenotype and a second may underlie the formation of neuronal synapses. We make cNMF available to the community and illustrate how this approach can provide key insights into gene expression variation within and between cell types.

## Introduction

Genes act in concert to maintain a cell’s identity as a specific cell type, to respond to external signals, and to carry out complex cellular activities such as replication and metabolism. Coordinating the necessary genes for these functions is frequently achieved through transcriptional co-regulation, where genes are induced together as a gene expression program (GEP) in response to the appropriate internal or external signal^1,2^. By enabling unbiased measurement of the whole transcriptome, profiling technologies such as RNA-Seq are paving the way for systematically discovering GEPs and shedding light on the biological mechanisms that they govern^3^.

Single-cell RNA-Seq (scRNA-Seq) has greatly enhanced our potential to resolve GEPs by making it possible to observe variation in gene expression over many individual cells. Even so, inferring GEPs remains challenging as scRNA-Seq data is noisy and highdimensional, requiring computational approaches to uncover the underlying patterns. In addition, technical artifacts such as doublets (where two or more distinct cells are mistakenly collapsed into one) can confound analysis. Methodological advances in dimensionality reduction, clustering, lineage trajectory tracing, and differential expression analysis have helped overcome some of these issues^4–7^.

Here, we focus on a key challenge of inferring expression programs from scRNA-Seq data: the fact that individual cells may express multiple GEPs but we only detect cellular expression profiles that reflect their combination, rather than the GEPs themselves. A cell’s gene expression is shaped by many factors including its cell type, its state in time-dependent processes such as the cell cycle, and its response to varied environmental stimuli^8^. We group these into two broad classes of expression programs that can be detectable in scRNA-Seq data: (1) GEPs that correspond to the identity of a specific cell type such as hepatocyte or melanocyte (identity programs) and (2) GEPs that are expressed independently of cell type, in any cell that is carrying out a specific activity such as cell division or immune cell activation (activity programs). In this formulation, identity programs are expressed uniquely in cells of a specific cell type, while activity programs may vary dynamically in cells of one or multiple types and may be continuous or discrete.

Thus far, the vast majority of scRNA-Seq studies have focused on systematically identifying and characterizing the expression programs of cell types composing a given tissue, i.e. identity GEPs. Substantially less progress has been made in identifying activity GEPs, primarily through direct manipulation of cells in controlled experiments, for example comparing stimulated and unstimulated neurons^9^ or cells pre- and post-viral infection^10^.

If a subset of cells profiled by scRNA-Seq expresses a given activity GEP, there is a potential to directly infer the program from the data without the need for controlled experiments. However, this can be significantly more challenging than ascertaining identity GEPs; while some cells may have expression profiles that are predominantly the output of an identity program, activity programs will always be expressed alongside the identity programs of one or frequently many cell types. Thus, while finding the average expression of clusters of similar cells may often be sufficient for finding reasonably accurate identity GEPs, it will often fail for activity GEPs.

We hypothesized that we could infer activity GEPs directly from variation in single cell expression profiles using matrix factorization. In this context, matrix factorization would model the gene expression data matrix as the product of two lower rank matrices, one encoding the relative contribution of each gene to each program, and a second specifying the proportions in which the programs are combined for each cell. We refer to the second matrix as a ‘usage’ matrix as it specifies how much each GEP is ‘used’ by each cell in the dataset. Unlike hard clustering, which reduces all cells in a cluster to a single shared GEP, matrix factorization allows cells to express multiple GEPs. Thus, this computational approach would allow cells to express one or more activity GEPs in addition to their expected cell-type GEP, and could correctly model doublets as a combination of the identity GEPs for the combined cell types. To the best of our knowledge, no previously reported studies have benchmarked the ability of matrix factorization methods to accurately learn identity and activity GEPs from scRNA-Seq profiles.

We see three primary motivations for jointly inferring identity and activity GEPs in scRNA-Seq data. First, systematic discovery of GEPs could reveal unexpected or novel activity programs reflecting important biological processes (e.g. immune activation or hypoxia) in the context of the native biological tissue. Second, it could enable characterization of the prevalence of each activity GEP across cell types in the tissue. Finally, accounting for activity programs could improve inference of identity programs by avoiding spurious inclusion of activity program genes in the latter. GEPs corresponding to different phases of the cell cycle are examples of widespread activity programs and are well-known to confound identity (cell type) program inference in scRNA-Seq data^11,12^. However, cell-cycle is just one instance of the broader problem of confounding of identity and activity programs.

While matrix factorization is widely used as a preprocessing step in scRNA-Seq analysis, a priori it is unclear which, if any, factorization approaches would be most appropriate for inferring biologically meaningful GEPs. In particular, Principal Component Analysis (PCA), Independent Component Analysis (ICA), Latent Dirichlet Allocation (LDA)^13^ and Non-Negative Matrix Factorization (NMF)^14^ have been used for dimensionality reduction of data prior to downstream analysis or as an approach to cell clustering. However, while PCA^10,15^, NMF^16^ and ICA^17^ components have been interpreted as activity programs, the dimensions inferred by these or other matrix factorization algorithms may not necessarily align with biologically meaningful gene expression programs and are frequently ignored in practice. This is because each method makes different simplifying assumptions that are potentially inappropriate for gene expression data. For example, NMF and LDA are non-negative and so cannot directly model repression. ICA components are statistically independent, PCA components are mutually orthogonal, and both allow gene expression to be negative. Furthermore, none of these methods, except LDA, explicitly accounts for the count distribution of expression data in their error models.

In this study, we motivate, validate, and enhance the use of matrix factorization for GEP inference. Using simulations, we show that despite their simplifying assumptions, ICA, LDA, and NMF--but not PCA--can accurately discover both activity and identity GEPs. However, due to inherent randomness in their algorithms, they give substantially varying results when repeated multiple times, which hinders their interpretability. We therefore implemented a meta-analysis approach (Fig. 1a), which demonstrably increased robustness and accuracy. Overall, the meta-analysis of NMF, which we call Consensus NMF (cNMF), gave the best performance in these simulations.

**Figure 1:**
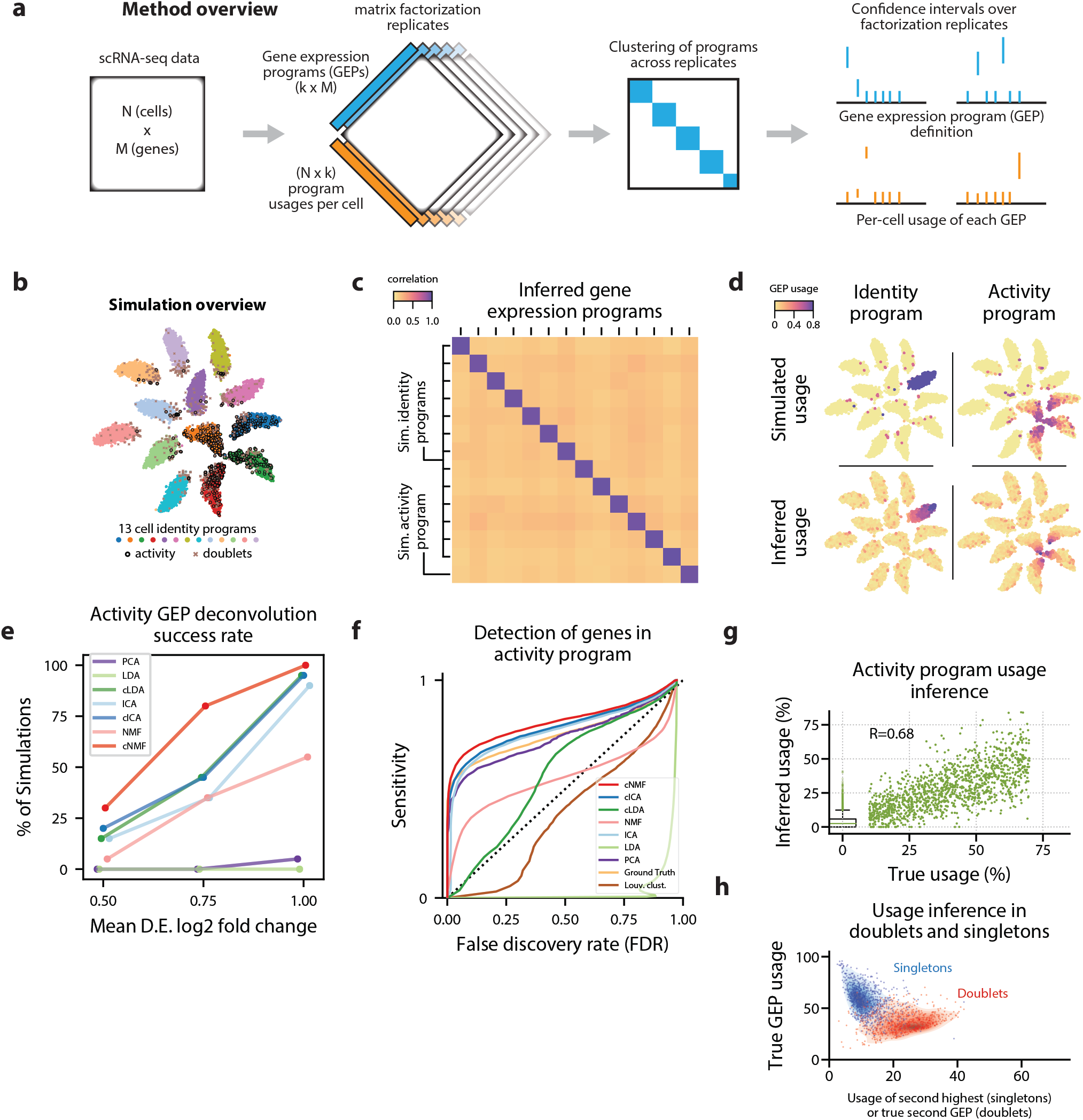
cNMF infers identity and activity expression programs in simulated data. **(a)** Schematic of the consensus matrix factorization pipeline. **(b)** t-distributed stochastic neigbour embedding (tSNE) plot of an example simulation showing different cell types with marker colors, doublets as gray Xs, and cells expressing the activity gene expression program (GEP) with a black edge. **(c)** Pearson correlation between the true GEPs and the GEPs inferred by cNMF for the simulation in (b). **(d)** Same tSNE plot as (b) but colored by the simulated or the cNMF inferred usage of an example identity program (left) or the activity program (right). **(e)** Percentage of 20 simulation replicates where an inferred GEP had Pearson correlation greater than 0.80 with the true activity program for each signal to noise ratio (parameterized by the mean log2 fold-change for a differentially expressed gene). **(f)** Receiver Operator Characteristic (except with false discovery rate rather than false positive rate) showing prediction accuracy of genes associated with the activity GEP. **(g)** Scatter plot comparing the simulated activity GEP usage and the usage inferred by cNMF for the simulation in (b). For cells with a simulated usage of 0, the inferred usage is shown as a box and whisker plot with the box corresponding to interquartile range and the whiskers corresponding to 5th and 95th percentiles. **(h)** Contour plot of the true GEP usage on the Y-axis and the second true GEP usage for doublets or the second highest GEP usage inferred by cNMF for singletons for the simulation in (b). 1000 randomly selected cells are overlayed as a scatter plot for each group.

Applied to three real datasets generated by 3 different scRNA-Seq platforms, cNMF inferred expected activity programs (cell-cycle programs in a brain organoid dataset and depolarization induced programs in visual cortex neurons), an unanticipated hypoxia program, and intriguing novel activity programs. It also enhanced cell type characterization and enabled estimation of rates of activity across cell types. These findings on real datasets further validate our approach as a useful analysis tool to understand complex signals within scRNA-Seq data.

## Results

### Evaluation of matrix factorization for GEP inference in simulated data

We sought to establish whether components inferred by simple matrix factorizations would align with GEPs in scRNA-seq data. We evaluated this in simulated data of 15,000 cells composed of 13 cell types, one cellular activity program that is active to varying extents in a subset of cells of four cell types, and a 6% doublet rate (Fig. 1b). We generated 20 replicates of this simulation, each at three different ‘signal to noise’ ratios, in order to determine how matrix factorization accuracy varies with noise level (Online methods).

We first analyzed the performance of ICA, LDA, and NMF and noticed that they yielded different solutions when run several times on the same input simulated data. We ran each method 200 times and assigned the components in each run to their most correlated ground-truth program. We saw that there was significant variability among the components assigned to the same program -- particularly for NMF and LDA (Supplementary Fig. 1). Unlike PCA, which has an exact solution, these factorizations use stochastic optimization algorithms to obtain approximate solutions in a solution space including many local optima. We observed that such local optima frequently corresponded to solutions where a simulated GEP was split into multiple inferred components and/or multiple GEPs were merged into a single component (Supplementary Fig. 2a). This variability reduces the interpretability of the solutions and may decrease the accuracy as well.

To overcome the issue of variability of solutions, we employed a meta-analysis approach, which we call consensus matrix factorization, that averages over multiple replicates to increase the robustness of the solution. The method which is adapted from a similar procedure in mutational signature discovery^18^ proceeds as follows: we run the factorization multiple times, filter outlier components (which tended to represent noise or merges/splits of GEPs), cluster the components over all replicates combined, and take the cluster medians as our consensus estimates. With these estimates fixed, we are able to compute a final usage matrix specifying the contribution of each GEP in each cell and to transform our GEP estimates from normalized units to biologically meaningful ones such as transcripts per million (TPM). This approach also provides us with a guide for choosing the number of components by selecting a value that provides a good trade-off between error and stability (Supplementary Fig. 3, Online methods). We refer to this approach as consensus matrix factorization based on its analogy with consensus clustering^19^ and to its application to LDA, NMF and ICA, as cLDA, cNMF, and cICA respectively (See materials and methods for details).

We found that the consensus matrix factorization approach inferred components underlying the GEPs as well as which cells expressed each GEP (Fig. 1c-d, Supplementary Figure 2, 4a). By contrast, principal components reflected linear combinations of the true GEPs. In addition to increasing the robustness, the consensus approach also increased the ability of factorization to deconvolute the true GEPs - most dramatically for LDA and NMF which had the most stochastic variability. cNMF successfully deconvoluted the activity and identity GEPs more frequently than the other matrix factorizations considered (Fig. 1e, Supplementary Fig. 3).

We next sought to benchmark the sensitivity and specificity of each matrix factorization method for inferring genes that are associated with each GEP. We also evaluated the performance of discrete clustering for this task because clustering is the most common way GEPs are identified in practice. We evaluated the commonly used Louvain community detection clustering algorithm^20,21^ but also considered an upper bound on how well any discrete clustering could perform by using ground-truth to assign cells to a cluster of its cell type or to an activity cluster if it had >=40% simulated contribution from the activity GEP (Supplementary Fig. 4b). We evaluated the association between genes and GEPs using linear regression and measured accuracy using a receiver operator characteristic (Online methods).

We found that cNMF was most accurate at inferring genes in the activity program, with a sensitivity of 61% at a false discovery rate (FDR) of 5% (Fig. 1f). cICA and the ground-truth clustering were the next most accurate with 57% and 56% sensitivity at a 5% FDR respectively. cNMF also performed the best at inferring identity GEPs of the 4 cell types that expressed the activity (Supplementary Fig. 5). As expected, the clustering approaches performed worse as they inappropriately assigned activity GEP genes to these identity programs, resulting in an elevated FDR. This illustrates how matrix factorization can outperform clustering for inference of the genes associated with activity and identity GEPs.

We decided to proceed with cNMF to analyze the real datasets due its accuracy, processing speed, and interpretability. First, it yielded the most accurate inferences in our simulated data. Second, NMF was the fastest of the basic factorization algorithms considered, which is especially useful given the need to run multiple replicates and given the growing sizes of scRNA-Seq datasets (Supplementary Fig. 6). Third, the nonnegativity assumption of NMF naturally results in usage and component matrices that can be normalized and interpreted as probability distributions--I.e. where the usage matrix reflects the probability of each GEP being used in each cell, and the component matrix reflects the probability of a specific transcript expressed in a GEP being a specific gene. The other high performing factorization method, cICA, produced negative values in the components and usages which precludes this interpretation.

Beyond identifying the activity program itself, we found that cNMF could also accurately infer which cells expressed the activity program and what proportion of their expression was derived from the activity program (Fig. 1g). With an expression usage threshold of 10%, cNMF accurately classified 91% of cells expressing the activity program and 94% of cells that did not express the program. Moreover, we observed a high Pearson correlation between the inferred and simulated usages in cells that expressed the program (R=0.74 for all simulations combined, R=0.68 for the example simulation in Fig. 1g). Thus, cNMF can be used both to infer which cells express the activity program, as well as what proportion of their transcripts derive from that program.

Finally, we demonstrated that cNMF was robust to the presence of doublets--instances where two cells are mistakenly labeled as a single cell. Due to limitations in the current tissue dissociation and single-cell sequencing technologies, some number of “cells” in an scRNA-Seq dataset will actually correspond to doublets. Several computational methods have been developed to identify cells that correspond to doublets but this is still an important artifact in scRNA-Seq data^22,23^. We found that cNMF correctly modeled doublets as a combination of the GEPs for the two combined cell types (Fig. 1h). Moreover, we found that cNMF could accurately infer the GEPs even in a simulated dataset composed of 50% doublets (Supplementary Fig. 7). This illustrates another benefit of representing cells in scRNA-Seq data as a mixture of GEPs rather than classifying them into discrete clusters.

### cNMF deconvolutes hypoxia and cell-cycle activity GEPs from identity GEPs in brain organoid data

Having demonstrated its performance and utility on simulated data, we then used cNMF to re-analyze a published scRNA-Seq dataset of 52,600 single cells isolated from human brain organoids^24^. The initial report of this data confirmed that organoids contain excitatory cell types homologous to those in the cerebral cortex and retina as well as unexpected cells of mesodermal lineage, but further resolution can be gained on the precise cell types and how they differentiate over time. As organoids contain many proliferating cell types, we sought to use this data to confirm that cNMF could detect activity programs -- in this case, cell cycles programs -- in real data, and to explore what biological insights could be gained from their identification.

We identified 31 distinct programs in this dataset that could be further parsed into identity and activity programs (Supplementary Fig. 8). We distinguished between identity and activity programs by using the fact that activity programs can occur in multiple diverse cell types while identity programs represent a single cell-type. Most cells had high usage of just a single GEP, which is consistent with expressing just an identity program (Fig. 2a). When cells expressed multiple GEPs, those typically had correlated expression profiles, suggesting that they correspond to identity programs of closely related cell types or cells transitioning between two developmental states, rather than activity programs (Supplementary Fig. 9). By contrast, 3 GEPs were co-expressed with many distinct and uncorrelated programs, suggesting that they represent activity programs that occur across diverse cell types (Fig. 2a-b). Consistent with this, the 28 suspected identity programs were well separated by the cell-type clusters reported in Quadrato et. al., while the three suspected activity programs were expressed by cells across multiple clusters (Supplementary Fig. 10-11). Except for a few specific cases discussed below, we used these published cluster labels to annotate our identity GEPs.

**Figure 2:**
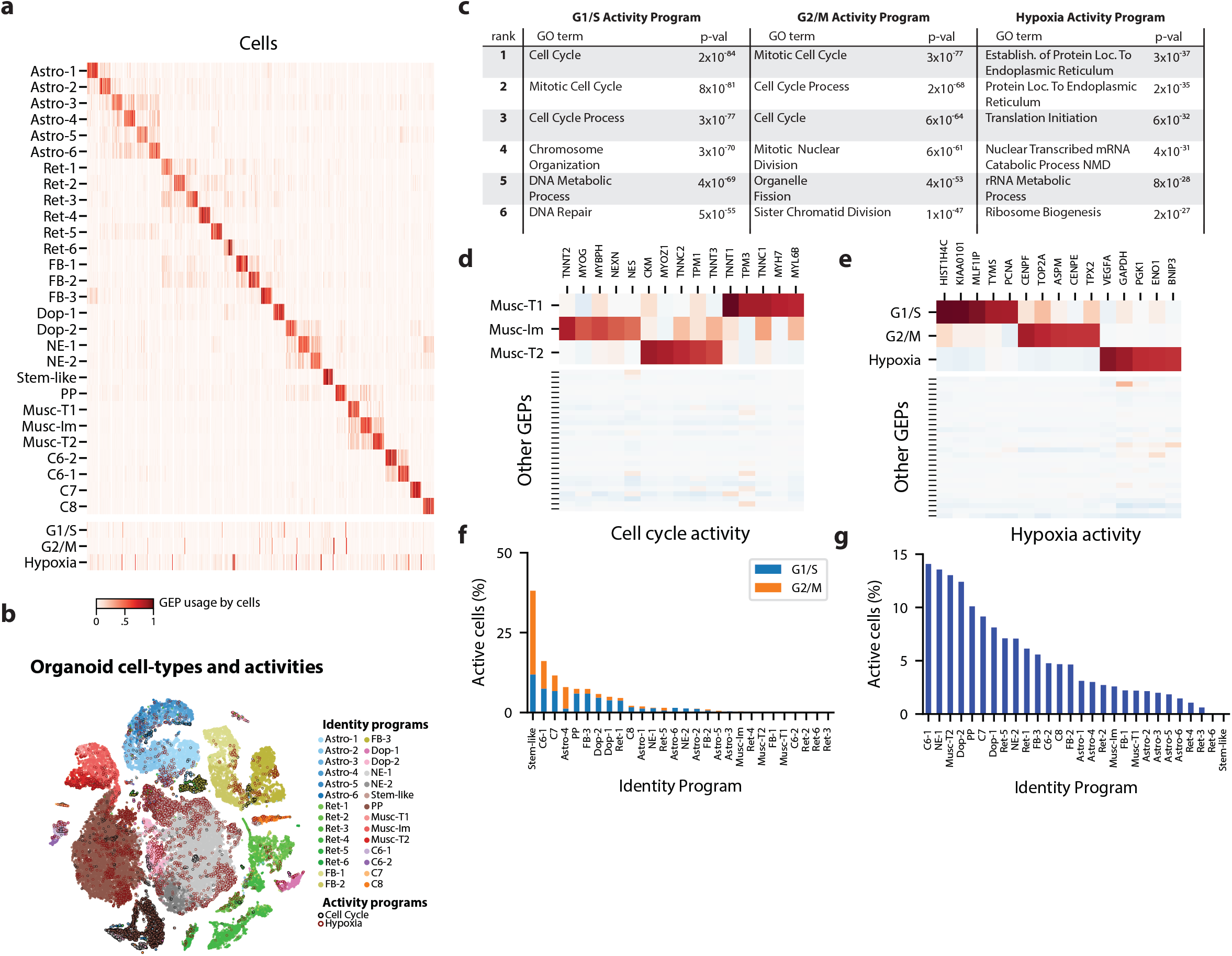
Deconvolution of activity programs from cell identity in brain organoid data. **(a)** Heat-map showing percent usage of all GEPs (rows) in all cells (columns). Identity GEPs are shown on top and activity GEPs are shown below. Cells are grouped by their maximum identity GEP and fit into columns of a fixed width for each identity GEP. **(b)** tSNE plot of the brain organoid dataset with cells colored by their maximally used identity GEP, and with a black edge for cells with >10% usage of the G1/S or G2/M activity GEP or a maroon edge for cells with >10% usage of the hypoxia GEP. **(c)**Table of P-values for the top six Gene Ontology geneset enrichments for the three activity GEPs. **(d)** Heatmap of Z-scores of top genes associated with three mesodermal programs, in those programs (top), and in all other programs (bottom). **(e)** Heatmap of Z-scores of top genes associated with three activity GEPs, in those programs (top), and in all other programs (bottom). **(f)** Proportion of cells assigned to each identity GEP that express the G1/S or G2/M program with a percent usage greater than 10%. **(g)** Proportion of cells assigned to each identity GEP that express the hypoxia program with a percent usage greater than 10%.

Our 28 identity programs further refined the 10 primary cell-type clusters originally reported for this dataset. For example, we noticed that cells previously annotated as mesodermal predominantly expressed one of three GEPs that were significantly enriched for genes in the ‘Muscle Contraction’ Gene Ontology (GO) set (P<1×10^−10^ vs. P>.19 for all other GEPs). They therefore likely represent muscle cells. Inspecting the genes associated with these 3 GEPs (Supplementary Table 1), we noticed that they include genes characteristic of different classes of skeletal muscle: (1) immature skeletal muscle (e.g. *MYOG, TNNT2, NES*), (2) fast-twitch muscle (e.g. *TNNT3, TNNC2, MYOZ1*), and (3) slow-twitch muscle (e.g. *TNNT1, TNNC1, TPM3*) (Fig. 2d). This unexpected finding suggests that distinct populations of skeletal muscle cells -- excitatory cell types with many similarities to neurons -- are differentiating in these brain organoids.

Of the three activity programs identified, we found that two were strongly enriched for cell cycle Gene Ontology (GO) sets, suggesting that they correspond to separate phases of the cell cycle (Fig. 2c). One showed stronger enrichment for genesets involved in DNA replication (e.g. DNA Replication P=3×10^−52^ compared to P=2×10^−3^) while the other showed stronger enrichment for genesets involved in mitosis (e.g. Mitotic Nuclear Division, P=4×10^−61^ compared to P=2×10^−46^). These enrichments and inspection of the genes most associated with these programs implied that one represents a G1/S checkpoint program and the other represents a G2/M checkpoint program (Fig. 2e). Thus, cNMF discovered two activity programs corresponding to separate phases of the cell cycle directly from the data.

The third activity program is characterized by high levels of hypoxia related genes (e.g. *VEGFA, PGK1, CA9, P4HA1, HILPDA*) suggesting it represents a hypoxia program (Fig. 2e). This is consistent with the lack of vasculature in organoids which makes hypoxia an important growth constraint^25^. This GEP is most significantly enriched for genesets related to protein localization to the endoplasmic reticulum and nonsense mediated decay (P=3×10^−37^, P=5×10^−31^) (Fig. 2c), consistent with reports that hypoxia post-transcriptionally increases expression of genes that are translated in the ER^26^ and modulates nonsense mediated decay activity^27^. In the initial report of this data, staining for a single hypoxia gene, *HIF1A*, failed to detect significant levels of hypoxia. Indeed, *HIF1A* is not strongly associated with this GEP, at least not at the transcriptional level. This highlights the ability of our unbiased approach to detect unanticipated activity programs in scRNA-Seq data.

Having identified proliferation and hypoxia activity programs, we sought to quantify their relative rates across cell types in the data. We found that 3079 cells (5.9%) expressed the G1/S program and 2043 cells (3.9%) expressed the G2/M program (with usage>=10%). Classifying cells into cell types according to their most used identity program, we found that many distinct populations were replicating. For example, cNMF detected a rare population, included with the forebrain cluster in the original report, that we label as “stem-like” based on high expression of pluripotency markers (e.g. *LIN28A, L1TD1, MIR302B, DNMT3B*) (Supplementary table 1). These cells showed the highest rates of proliferation with over 38% of them expressing a cell-cycle program in addition to the “stem-like” identity GEP (Fig. 2f).

cNMF was further able to refine cell types by disentangling the contributions of identity and activity programs to the gene expression of cells. For example, we found that a cell cluster labeled in Quadrato et al., 2017 as “proliferative precursors”, based on high expression of cell-cycle genes, is composed of multiple cell types including immature muscle and dopaminergic neurons (Supplementary Fig. 11). The predominant identity GEP of cells in this cluster is most strongly associated with the gene PAX7, a marker of self-renewing muscle stem cells^28^ (Supplementary table 1). Indeed, this GEP has high (>10%) usage in 41% of cells who’s most used GEP is the immature muscle program, suggesting it may be a precursor of muscle cells. This relationship was not readily identifiable by clustering because the majority of genes associated with the cluster were cell cycle related.

We also saw a wide range of cell types expressing the hypoxia program, with the highest rates in C6-1, neuroepithelial-1, type 2 muscle, and dopaminergic-2 cell types. The lowest levels of hypoxia program usage occurred in forebrain, astroglial, retinal, and type 1 muscle cell types (Fig. 2g). The hypoxia response program is widespread in this dataset with 5,788 cells (11%) of all cells expressing it (usage > 10%). This illustrates how inferring activity programs in scRNA-Seq data using cNMF makes it possible to compare the rates of cellular activities across cell types.

### cNMF identifies depolarization induced and novel activity programs in scRNA-Seq of mouse visual cortex neurons

Next we turned to another published dataset to further validate cNMF and to illustrate how it can be combined with scRNA-Seq of experimentally manipulated cells to uncover more subtle activity programs. We re-analyzed scRNA-Seq data from 15,011 excitatory pyramidal neurons or inhibitory interneurons from the visual cortex of dark-reared mice that were suddenly exposed to 0 hours, 1 hours, or 4 hours of light^9^. This allowed the authors to identify transcriptional changes induced by repeated depolarization, a phenomenon believed to be critical for proper cortical function. We sought to determine whether cNMF would identify the relatively modest activity programs (~60 genes with fold-change >=2 and FDR<0.05) elicited by the experiment without knowledge of the experimental design labels. Furthermore, since the authors identified heterogeneity in stimulus-responsive genes between neuronal subtypes, we wondered if cNMF would identify a common activity program and whether it could tease out patterns in what is shared or divergent across neuron subtypes.

We ran cNMF on neurons combined from all three exposure conditions and identified 20 GEPs, interpreting 14 as identity and 6 as activity programs (Supplementary Fig. 12). As we saw in the organoid data, the activity programs were co-expressed with many distinct and uncorrelated GEPs while the identity programs only overlapped in related cell types (Fig. 3a-b). In addition, the identity programs were well separated by the published clusters while the activity programs were spread across multiple clusters (Supplementary Fig. 13). We thus used the published cluster annotations to label the identity GEPs.

**Figure 3:**
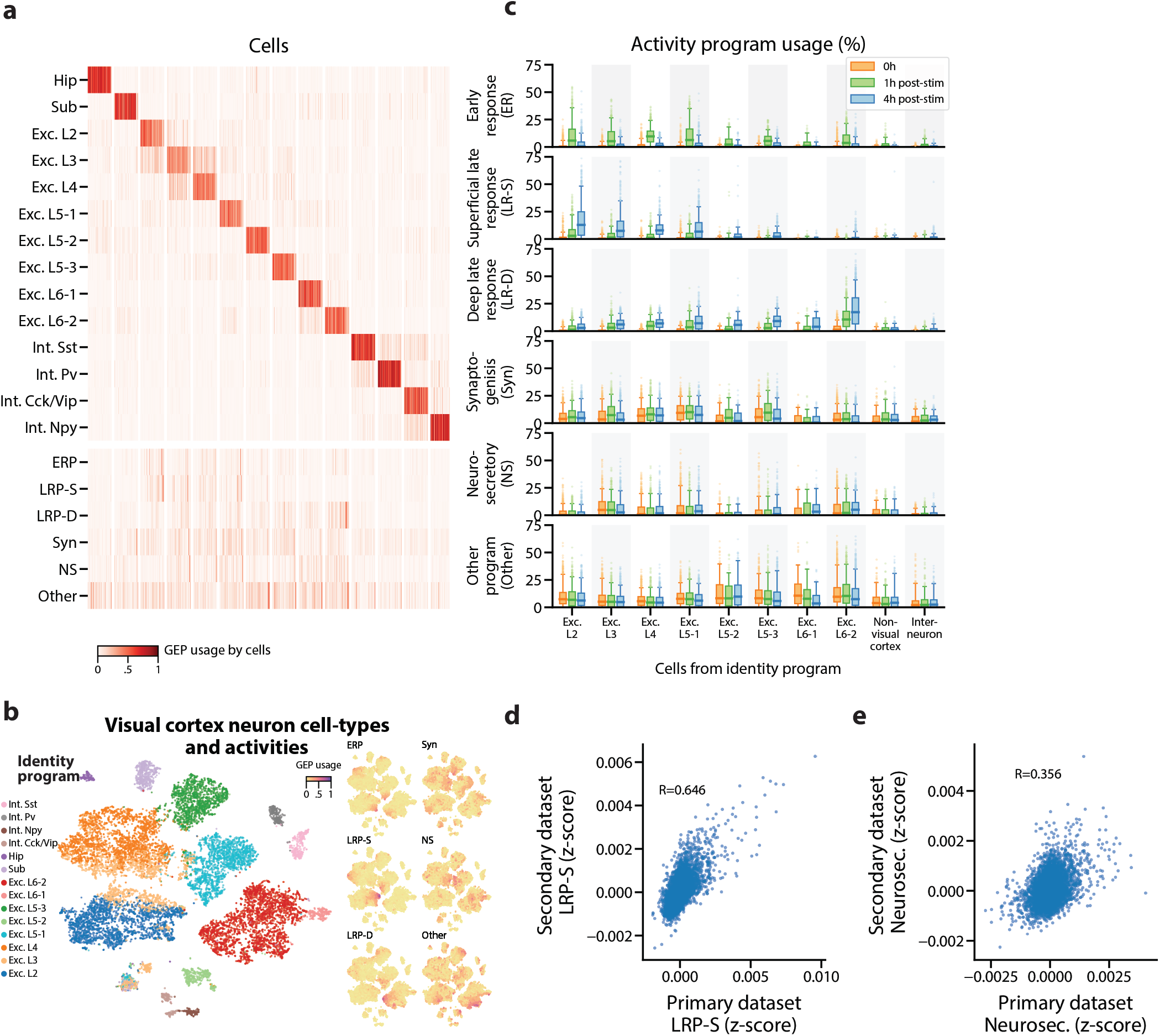
Identification of activity programs in neurons of the visual cortex. **(a)** Heatmap showing percent usage of all GEPs (rows) in all cells (columns). Identity GEPs are shown on top and activity GEPs are shown below. Cells are grouped by their maximum identity GEP and fit into columns of a fixed width for each identity GEP. **(b)** t-SNE plots of cells colored by maximum identity GEP usage (left) or by absolute usage of each activity GEP (right). **(c)** Box and whisker plot showing the percent usage of activity programs (rows) in cells classified according to their maximum identity GEP (columns) and stratified by the stimulus condition of the cells (hue). The central line represents the median, boxes represent the interquartile range, and whiskers represent the 5th and 95th quantile. **(d)** Scatter plot of Z-scores of the superficial late response GEP in the primary dataset against the corresponding GEP in the Tasic et. al., 2016 dataset. **(e)** Same as (d) but for the neurosecretory activity program.

Three activity programs were correlated with the stimulus, which indicates that they are induced by repeated depolarization (Fig. 3c). One of these was induced at 1H and thus corresponds to an early response program (ERP). The others were primarily induced at 4H and thus correspond to late response programs (LRPs). These programs overlapped significantly with the sets of differentially expressed genes reported in Hrvatin et. al. 2017 (P=8×10^−34^ for the ERP and genes induced at 1H; P=4×10^−22^, P=4×10^−14^ for the LRPs and genes induced at 4Hs, one-sided Mann Whitney test).

Intriguingly, one LRP was more induced in superficial cortical layers, while the other was more induced in deeper layers. This supports a recently proposed model where the ERP is predominantly shared across excitatory neurons, while LRPs vary more substantially across neuron subtypes^9^. It also illustrates cNMF’s sensitivity: in the initial report, only 64 and 53 genes were identified as differentially expressed in at least one excitatory cell type at 1H and 4Hs (FC≥2, FDR<.05). Nevertheless, cNMF was able to find this program in an unsupervised manner, without knowledge of the experimental design.

cNMF was also able to identify a depolarization induced program in visual cortex neurons that were not experimentally manipulated to elicit them. We analyzed an additional scRNA-Seq dataset of 1,573 neurons from the visual cortex of adult mice that, unlike in the primary dataset, were not reared in darkness or treated with a specific light stimulus^29^. In this dataset, cNMF identified a matching GEP for all visual cortex cell types found in the primary dataset except for a single subtype of excitatory layer 5 (Supplementary Fig. 14a). Moreover, it identified a GEP that showed striking concordance with the superficial LRP found in the primary dataset (Fisher Exact Test of genes with association Z-score>.0015, OR=127, P=1×10^−118^, Pearson Correlation=.645) (Fig. 3d). This program was predominantly expressed in excitatory cells of the more superficial layers of the cortex as would be expected based on the results in the primary dataset. For example, over 40% of the excitatory layer 2 (Exc. L2) type neurons expressed this activity program (Supplementary Fig. 14b). This demonstrates that cNMF could also find the depolarization induced activity program in scRNA-Seq of cells that had not been experimentally manipulated to elicit it.

Finally, cNMF identified three additional activity programs in the primary visual cortex dataset that were not well correlated with the light stimulus but were expressed broadly across multiple neuronal cell types (Fig. 3b-c). We labeled one of these, that was specific to excitatory neurons, as Neurosecretory (NS) because it is characterized by high expression of several secreted neuropeptides including *Vgf, Adcyap1, Scg2, Cck, Scg3*, and *Dkk3*, and has high expression of genes that facilitate protein secretion such as *Cpe, Cadps2*, and *Scamp5* (Supplementary table 2). The top expressed gene--*Vgf* (VGF nerve growth factor inducible) is induced by nerve growth factor^30^, suggesting that this program may be regulated by external growth factor signals. Notably, we found a matching program in the Tasic Et, Al. dataset (Fisher Exact Test of genes with association Z-score>.0015, OR=53.8, P=8×10^−21^, Pearson Correlation=.356) (Fig. 3e). Thus, this neurosecretory activity program is reproducible across multiple single-cell datasets.

An additional activity program which we labeled Synaptogenesis (Syn) was characterized by expression of genes that play a crucial role in synapse formation, including the transcriptional regulator *Mef2c^31–33^*, synaptic adhesion molecules *Ncam1^34^* and *Cadm1^35,36^*, membrane vesicle traffickers *Syt1* and *Syt4, Actb* which constitutes the predominant synapse cytoskeletal protein, and others with a strong connection to synapse biology such as *Ywhaz*^37–39^, *Bicd1^40,41^*. It was also significantly enriched for relevant Gene Ontology sets including postsynapse, glutamatergic synapse, postsynaptic density, and dendrite morphogenesis (P<=3.25×10^−6^ Supplementary Table 3) which further suggests its interpretation as a program involved in the formation or regulation of synapses. The last activity program (labeled Other) was characterized by high expression of the maternally expressed long non-coding RNA *Meg3^42^* and other genes that are associated with cerebral ischemic injury (e.g. *Glg1^43^, Rtn1^44^*). Our functional interpretations of the novel activity programs are speculative, but they highlight the ability of cNMF to identify intriguing GEPs in an unbiased fashion.

## Discussion

scRNA-Seq is enabling discovery of the repertoire of GEPs that underlie the behaviors of cells in complex organisms. Here, we distinguish between cell type (identity) and cell type independent (activity) expression programs to describe the types of GEPs that can be inferred with the resolution of current scRNA-Seq datasets. More subtle GEPs, such as those that vary based on stochastic fluctuations in individual transcription factors, may eventually become discernible from scRNA-Seq data but currently remain outside the reach of current datasets. We note that some biological programs might blur the line between identity and activity GEPs, including programs reflecting oncogenic transformation, or a cell’s position along a morphological gradient. These examples represent continuums of transcriptional states that blur the line between cell type and cellular activity. Nevertheless, matrix factorization should, in theory, be able to capture these biological phenomena as long as the transitions between expression states can be modeled as linear combinations of the contributing GEPs. This may be a reasonable approximation for many biological transitions.

In this work, we have provided an empirical foundation for the use of matrix factorization to simultaneously infer identity and activity programs from scRNA-Seq data. We first show with simulations that despite their simplifying assumptions, ICA, LDA, and NMF (but not PCA) can infer components that align well with GEPs. However, due to the stochastic nature of these algorithms, the interpretability and accuracy of individual solutions can be low. This led us to develop a consensus approach that empirically increased the accuracy and robustness of the solutions. cNMF inferred the most accurate identity and activity programs of all the methods we tested. Moreover, it yielded results in interpretable units of gene expression (transcripts per million) and could accurately infer the percentage of each cell’s expression that was derived from each GEP. These properties made it the most promising approach for GEP inference on real datasets.

We explored the utility of cNMF on real data, recapitulating known GEPs, identifying novel ones, and further characterizing their usage. We first validated cNMF with several expected activity programs serving as positive controls. We then identified several unexpected but highly plausible programs, a hypoxia program in brain organoids and a depolarization induced activity program in untreated neurons. Finally, we identified three novel programs in visual cortex neurons that we speculate may correspond to a neurosecratory phenotype, new synapse formation, and a stress response program. Beyond simply discovering activity programs, cNMF clarified the underlying cell types in these datasets by disentangling activity and identity programs from the mixed single-cell profiles. For example, we found that a brain organoid subpopulation that was initially annotated as proliferative precursors actually includes replicating cells of several cell types, including an immature skeletal muscle cell that is differentiating into slow-twitch and fast twitch muscle populations. Furthermore, joint analysis of identity and activity GEPs allowed us to quantify the relative prevalence of activities across cell types. For example, we found in the visual cortex data that one depolarization-induced late response program was predominantly expressed in neurons of superficial cortical layers, while the other was mainly expressed in deeper layers. This suggests that an anatomical or developmental factor may underlie variability in the response. While commonly used approaches based on clustering or pseudotemporal ordering of cells are poorly suited to achieve such insights from single-cell data, these findings emerge naturally from our matrix factorization approach.

We have made our tools and analyses readily accessible so that researchers can readily use cNMF and further develop on the approach. We have deposited all of the cNMF code on Github https://github.com/dylkot/cNMF/ and have made available all of the analysis scripts for figures contained in this manuscript on Code Ocean (https://doi.org/10.24433/CO.9044782e-cb96-4733-8a4f-bf42c21399e6) for easy exploration and re-execution.

As others apply this approach, one key consideration will be the choice of the 3 input parameters required by cNMF: the number of components to be found (K), the percentage of replicates to use as nearest neighbors for outlier-detection, and a distance threshold for defining outliers. While the choice of K must ultimately reflect the resolution desired by the analyst, we propose a simple decision aid of considering the trade-off between factorization stability and reconstruction error (Supplementary figures 3, 7, 8, 12). In addition, we noticed that choosing consecutive values of K primarily influenced individual components at the margin, suggesting that cNMF may be robust to this choice within a range of several options. The additional parameters allow users to optionally identify outlier replicates to filter before averaging across replicates. This improves overall accuracy by removing infrequent solutions that often represent merges or splits of the true GEPs. Using 30-35% of replicates as nearest neighbors worked well for all datasets we analyzed, and an appropriate outlier distance threshold was clear in our applications based on the long tail in the distance distribution (Supplementary figures 3, 7, 8, 12).

Our approach is an initial step forward towards disentangling identity and activity GEPs, and will benefit from subsequent development. For example, cNMF does not specifically address the count nature of gene expression nor the possibility of dropout events in its error model. Recently developed statistical frameworks that address these aspects of scRNA-Seq data such as Zero-inflated Factor Analysis^45^ and Hierarchical Poisson Factorization^46^ may therefore increase the accuracy of GEP inference. In addition, NMF often yields low but non-zero usages for many GEPs even though we expect most cells to express a small number of identity and activity GEPs. This lack of sparsity is likely due to over-fitting and could be addressed by adding regularization to the model^47^. Such refinements and any new matrix factorization that relies on stochastic optimization can be readily combined with our consensus approach to potentially improve accuracy and interpretability.

A more fundamental limitation of matrix factorizations, including cNMF, is the built-in assumption that cells can be modeled as linear combinations of GEPs. Notably, this precludes modeling of transcriptional repression, where one or more genes that would be induced by one GEP are significantly reduced in expression when a second repressing GEP is active in the same cell. To our knowledge such relationships have not been represented in a matrix factorization framework, but they may be easier to incorporate in new classes of latent variable models such as variational auto-encoders (VAEs)^48,49^. VAEs represent cells in a highly flexible latent space that can capture nonlinearities and interactions between latent variables. However, while the latent variables are designed to facilitate accurate reconstruction of the input gene expression data, it remains to be shown whether they can be directly or indirectly interpreted as distinct GEPs and GEP usages. For the foreseeable future, there may be a trade-off between the flexibility of these models and the difficulty in training them and interpreting their output.

With ongoing technological progress in RNA capture efficiency and throughput, scRNA-Seq data is likely to become richer and more expansive. This will make it possible to detect increasingly subtle GEPs, reflecting biological variability in cell types, cell states, and activities. Here, we have demonstrated a computational framework that can be used to infer such GEPs directly from the scRNA-Seq data without the need for experimental manipulations, providing key insights into the behavior of cells and tissues.

## Acknowledgements

We thank Allon Klein, Samuel Wolock, Aubrey Faust, Chris Edwards, Stephen Schaffner, the CGTA discussion group, and members of the Sabeti Laboratory for useful discussions and feedback on the manuscript. We thank the Arlotta, Greenberg, and Zeng laboratories for generating the primary datasets we analyze in this manuscript. The project described was supported by award Number R01AI099210 from the National Institute of Allergy and Infectious Disease and T32GM007753 from the National Institute of General Medical Sciences. The content is solely the responsibility of the authors and does not necessarily represent the official views of the National Institute of General Medical Sciences or the National Institutes of Health.

## Author contributions

DK and AV conceived of the project, developed the method, analyzed the data, and wrote the manuscript with input from the other authors.

MAN provided crucial guidance in analyzing the visual cortex data.

ST helped with implementing early versions of the method.

EH, DAM, and PCS provided guidance on the project.

## Materials and methods

### Simulations

Our simulation framework is based on Splatter^50^ but is re-implemented in Python and adapted to allow simulation of doublets and activity programs. Gene-expression programs were simulated as in Splatter. Cells were then randomly assigned an identity program with uniform proportions. 30% of cells of 4 cell types were randomly selected to express the shared activity program at a usage that was uniformly distributed between 10% and 70%. Each cell’s mean gene-expression profile was computed as the weighted sum of their cell-identity program and the activity program. Doublets were constructed by randomly sampling pairs of cells, summing their gene counts, and then randomly down-sampling the counts to the maximum of the two cells. We simulated 25,000 genes, 1,000 of which were associated with the activity program. The probability of a gene being differentially expressed in a cell identity program was set to 2.5%. The differential expression scale parameter was 1.0 for all simulations and the location parameter was either 1.0, 0.75, or 0.5 to simulate different signal to noise levels. Other splatter parameters were: lib.loc=7.64, libscale=0.78, mean_rate=7.68, mean_shape=0.34, expoutprob=0.00286, expoutloc=6.15, expoutscale=0.49, diffexpdownprob=0, bcv_dispersion=0.448, bcv_dof=22.087. These values were inferred from 8000 randomly sampled cells of the Quadrato et al., 2017 organoid dataset using Splatter. The same parameters and a differential expression location parameter of 1.0 were used for the 50% doublet simulation (Supplementary Figure 7).

### Data preprocessing

For each dataset, we removed cells with fewer than 1000 unique molecular identifiers (UMIs) detected. We also filtered out genes that were not detected in at least 1 out of 500 cells. We then selected the 2000 genes with the most over-dispersion as determined by the v-score^51^ for input to cNMF. Each high-variance gene was scaled to unit variance before running cNMF. This is similar to the log transformation that is commonly applied to scRNA-Seq data in that it ensures genes on different expression scales contribute comparable amounts of information to the programs. However, this avoids the need for addition of a pseudocount or modulation of the shape of a gene’s distribution. We do not mean center the genes so as to preserve the non-negativity of the expression data, which is a requirement for NMF.

Note, we do not perform any cell count normalization prior to cNMF. This is because cells with more counts can contribute more information to the model. Technical variation in transcript abundances across cells are captured in the Usage matrix rather than the component matrix. However, for the Tasic Et. Al. dataset, which is based on full-transcript sequencing rather than digital UMI counting, we variance-normalized high-variance genes from the TPM matrix directly rather than from the raw count matrix as in the other datasets.

As a final step in cNMF, the consensus programs can be re-fit including all genes and scaled to meaningful biological units of the user’s choice, such as transcripts per million (TPM) (see below).

### Consensus Non-negative Matrix Factorization (cNMF)

We use non-negative matrix factorization implemented in scikit-learn (version 20.0) with the default parameters except for random initialization, tolerance for the stopping condition of 10^−4^, and a maximum number of iterations of 400.

Each replicate of cNMF is run with a randomly selected seed and the same normalized dataset consisting of high-variance genes. The component matrices from each replicate are concatenated into a single matrix where each row is an NMF component from one replicate. Each of these components is normalized to have L2 norm of 1. Then components with high average euclidean distance from their K nearest neighbors are filtered out. We set the number of nearest neighbors, K, to be .3 times the number of bootstraps for the organoid and visual cortex datasets and .35 times the number of bootstraps for the simulated datasets. The threshold on average euclidean distance to nearest neighbors was set by inspecting the histogram and truncating the long tail (Supplementary Figs. 3, 7, 8, 12). Then, the replicate components were clustered using KMeans with the Euclidean distance metric and the same number of clusters as the number of components for the NMF runs. Each cluster is then collapsed down to a single component by taking the median value for each gene across components in a cluster. These merged GEP components are then normalized to sum to 1 and a final usage matrix is fit by running one last iteration of NMF with the component matrix fixed to this value. With this usage matrix fixed, final program estimates can be computed in desired units, and for all genes--including ones that were not initially included among the over-dispersed set. This is done by running a last iteration of NMF with the usage matrix fixed and the input data reflecting the desired final units. To convert the estimated programs to TPM scale, we refit against the TPM matrix.

We guide the selection of the number of cNMF components using the approach described in Alexandrov et al, 2013^18^ with a few modifications. We run NMF on normalized data matrices rather than count matrices and therefore do not resample counts but simply repeat NMF with different randomly selected seeds. We still guide determination of the number of components by considering the trade-off between Frobenius error and silhouette score of the clustering. However, we used the Frobenius error of the consensus NMF solution but without any outlier filtering. As in Alexandrov et al, 2013, stability is computed as the Silhouette score of the KMeans clustering on the NMF components (prior to filtering outliers). However we use Euclidean distance on L2 normalized components as the metric rather than Cosine distance. Silhouette score is calculated using the Scikit-learn version 20.0 silhouette_score function.

We parallelized the individual factorization steps over cores on a multi-core virtual machine using GNU Parallel^52^.

### Finding genes associated with programs

We identify genes that have specifically higher or lower expression in a GEP relative to background using Ordinary Least Squares Regression. We use the consensus usage matrix (cells x programs) for each cell as the predictor. For discrete clustering methods, used in the comparison, the usage matrix is a binary indicator matrix containing a 1 for the cluster (column) each cell is assigned to, and a 0 for all other columns. We regress this usage matrix against the TPM profile for each gene, normalized to mean 0 and variance 1. Thus, a positive coefficient for a GEP indicates that cells with a higher usage of the GEP will have higher expression of the gene than average, all else equal. The regression coefficient can be interpreted as signifying by how many standard deviations a cell’s expression would be expected to change from an additional count attributed to a GEP.

### Comparison of cNMF with other methods

We compared cNMF with consensus and standard versions of LDA and ICA as well as with PCA, Louvain clustering and a clustering based on assignment of cells to their ground-truth labels. We used the implementations of LDA, ICA, and PCA in scikit-learn and the implementation of Louvain clustering in scanpy^53^. For ICA, we used the FastICA implementation with default options for all the parameters. For LDA, we used the batch algorithm and all other parameters as defaults. We defined the consensus estimates across 200 replicates in the same way as for cNMF but with a slight modification for ICA. Because elements of the ICA components can either be positive or negative, some iterations would produce a components pointed in one direction while others would produce approximately the same component but pointed in the opposite direction (multiplied by −1). Therefore, we aligned the orientation of components from across replicates by identifying components whose median usage across all cells was positive and scaled these and the corresponding usages by −1.

For Louvain clustering, we used 14 principal components to compute distances between cells and used 15 nearest neighbors to define the KNN graph. For ground-truth assignment clustering, we assigned each cell to a cluster defined by its true identity program, except for cells which had greater that 40% usage of the activity program, which we assigned to an activity program cluster. Then we determined a GEP corresponding to each cluster as the mean TPM value for each gene over cells in the cluster.

To evaluate the accuracy of these various methods, we first calculated the coefficient for associating each genes with each program as described above. We then calculated sensitivity and false discovery rate (FDR) for each threshold on those coefficients and plotted those as an ROC-curve, except with FDR on the X-axis instead of false positive rate. For this evaluation, we considered a gene as truly associated with a GEP if it had a ground-truth fold-change of >=2 and only considered genes as null if the had no fold-change.

### Testing enrichment of genesets in programs

We used the Z-score regression coefficients identified as above, but with negative coefficients floored to 0, as inputs for a one-sided Mann Whitney U Test (with tie correction) comparing the median of genes in each geneset to those of genes not in the geneset.

### Data Availability

The organoid data described in the manuscript is accessible at NCBI GEO accession number GSE86153. However, we obtained the clustering and unnormalized data by request from the authors. The visual cortex datasets used for Figure 3 are accessible at NCBI GEO, accession numbers GSE102827 and GSE71585. All datasets (including the simulation) and the code to reproduce all analyses in this manuscript are available through Code Ocean (link: TBD)

### Code Availability

Code for running cNMF is available on Github https://github.com/dylkot/cNMF, as is code for simulating data with doublets and activity programs https://github.com/dylkot/scsim.

All analysis in this manuscript are available for exploration and re-execution on Code Ocean: https://doi.org/10.24433/CO.9044782e-cb96-4733-8a4f-bf42c21399e6

---

**Supplementary Figure 1:**
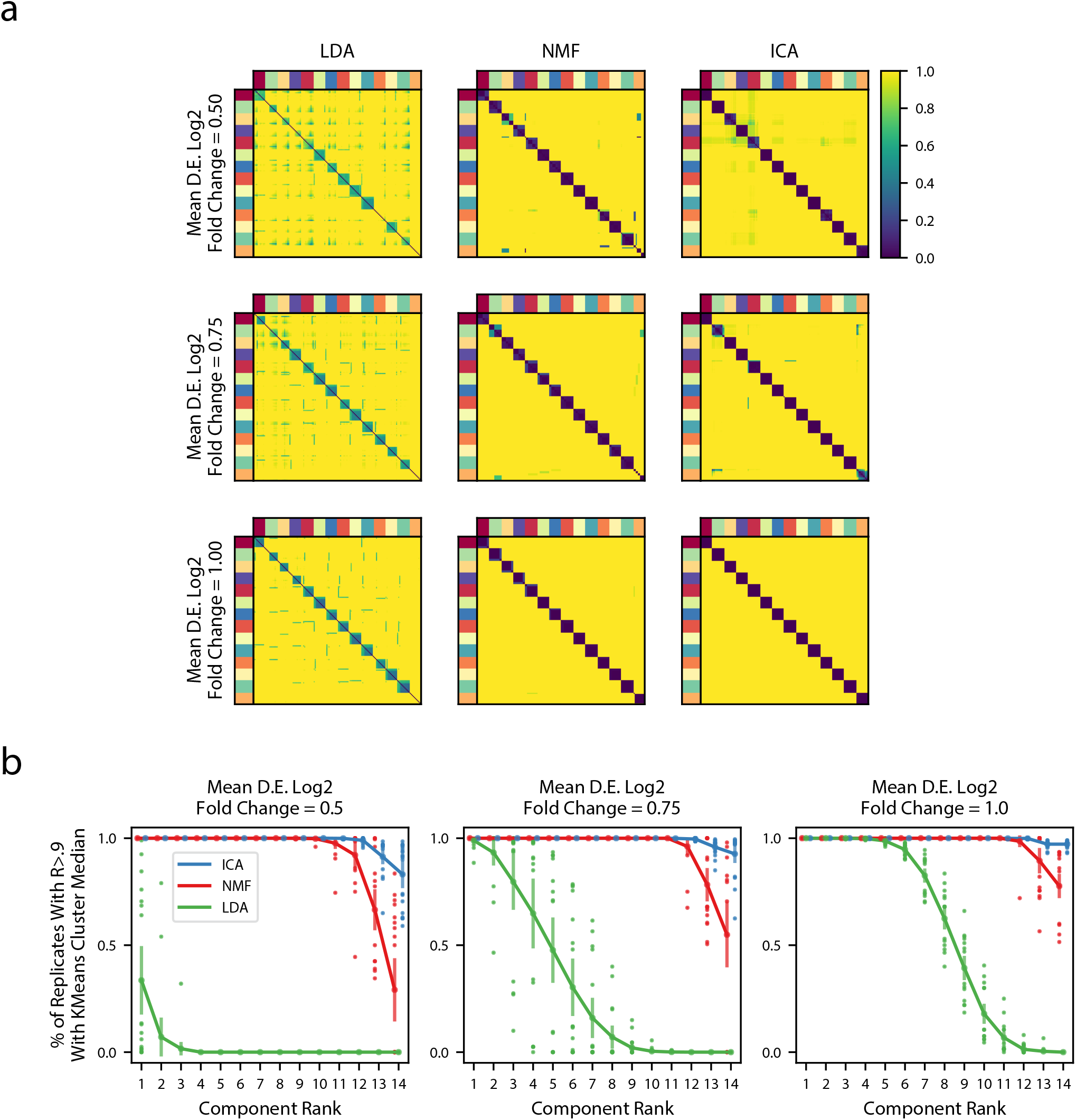
Robustness of Matrix Factorization Methods. **(a)** Pairwise correlation coefficients for all components combined across 200 replicates for multiple simulations (rows) and multiple matrix factorization algorithms (columns). All 14 components for each replicate were assigned in a one-to-one fashion to a ground truth GEP in order from most correlated to least correlated. Components are ordered in the plot by the assigned ground truth GEP which is denoted by the outer color bars. The three simulations had different average signal strengths as parameterized by the mean log2 fold-change for differentially expressed genes in a GEP (Online Methods) and are ordered from least signal (top row) to most signal (bottom row) **(b)** Components from the 200 replicates were assigned to a ground truth GEP, as in (a), and were correlated with the median of their assigned group. Then, for each factorization replicate, the 14 components were ranked in order from most to least correlated component. The plots shows the % of replicates where the kth most correlated component had a Pearson correlation >.9 with the median of the components assigned to the same ground truth GEP. We plot the mean and standard deviation of this percentage across the 20 replicates as bars around the mean, and plot individual dots for simulation replicates that exceeded one standard deviation of the mean.

**Supplementary Figure 2:**
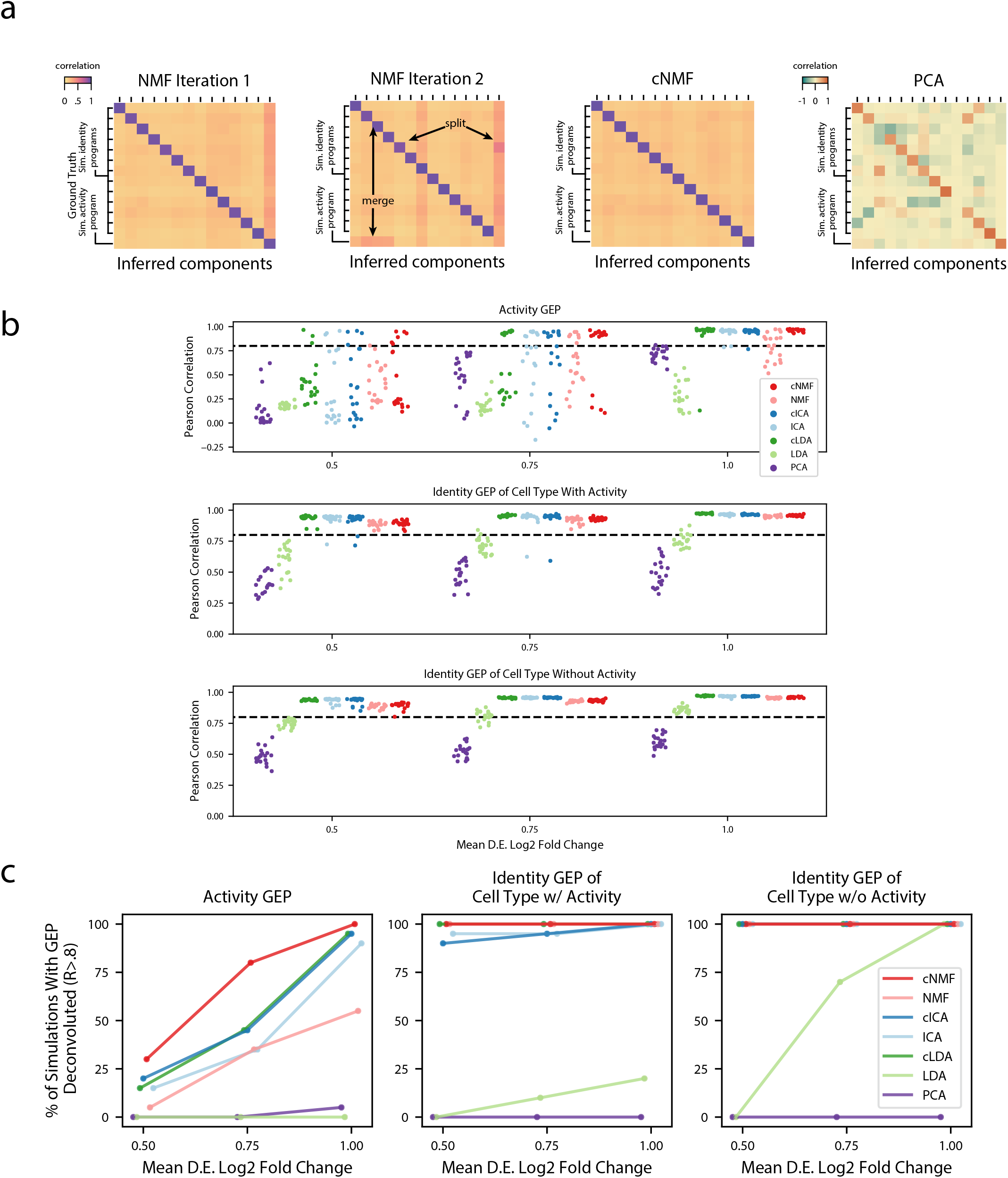
Deconvolution accuracy of matrix factorization methods. **(a)** Pearson correlation between ground truth GEP means (rows) and GEPs inferred by different iterations of NMF (left), cNMF, or PCA (right) for an example simulation. All correlations are computed considering only the 2000 most over-dispersed genes and on vectors normalized by the computed sample standard deviation of each gene. Arrows annotate cases where GEPs were merged into a single component or a GEP was split into 2 components. **(b)** Pearson correlation between inferred GEPs and true simulated GEPs for several matrix factorization methods across all 20 simulation replicates for each average signal level. **(c)** Percentage of simulation replicates for which GEPs of each type had a pearson correlation of R>.8 with the true simulated GEP, as a function of the average signal level.

**Supplementary Figure 3:**
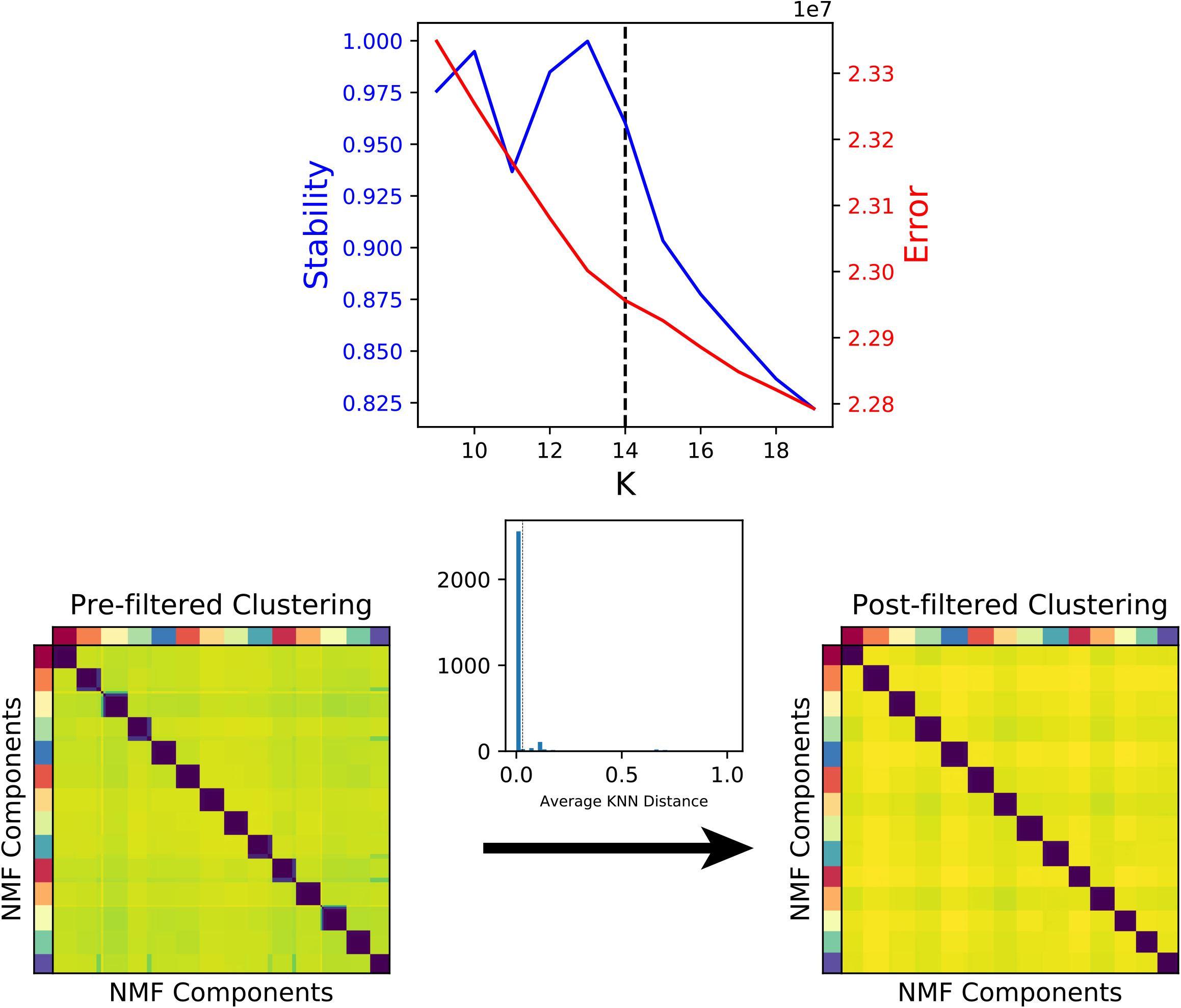
Diagnostic plot for cNMF on an example simulated dataset. **(a)** Number of cNMF components (K) against solution stability (blue, left axis) measured by the euclidean distance silhouette score of the clustering, and Frobenius error of the consensus solution (red, right axis). **(b)** Clustergram showing the clustered NMF components for K=14, combined across 200 replicates, before (left) and after filtering (right). In between, we show the average distance of each component to its 70 nearest neighbors as a histogram with a dashed line where we set the threshold for filtering outliers.

**Supplementary Figure 4:**
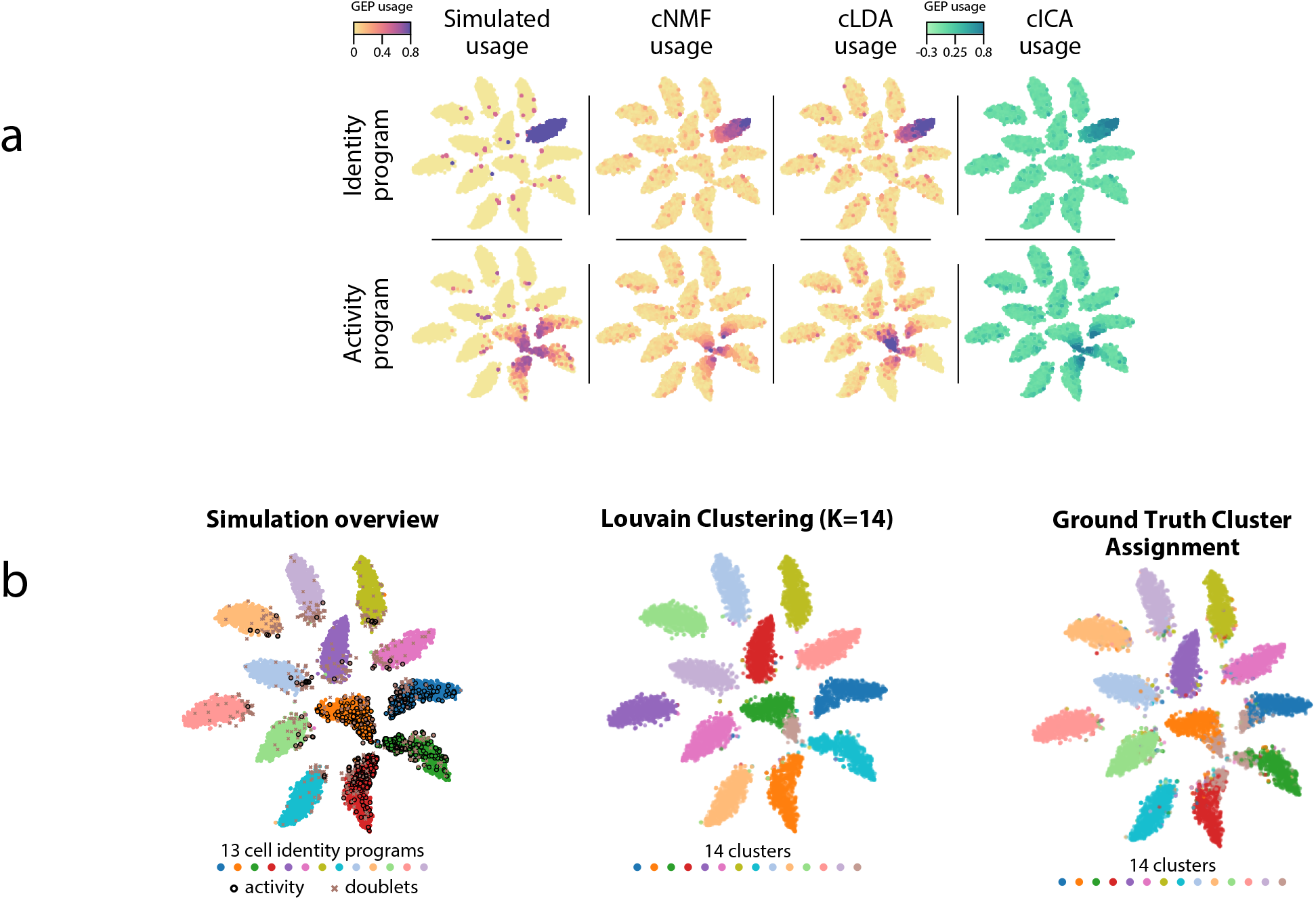
GEP usage inference. **(a)** Comparison of GEP usage inference by cNMF, cLDA, and cICA for an example identity GEP (top row) and the activity GEP (bottom row). Each cell is represented as a point and its usage is represented by the marker color. **(b)** Comparison of the results of ground truth cluster assignment and Louvain clustering represented on a t-SNE plot. (Left) Reproduction of figure 1b which shows cells colored based on their true identity program, doublets marked with an X, and cells that express the activity program with a black border. (Middle) Cells are colored based on Louvain clustering. (Right) Cells with activity GEP usage of greater than 40% are assigned to an activity cluster, and all other cells are assigned to their identity cluster. This shows how an ‘optimal’ discrete clustering might behave.

**Supplementary Figure 5:**
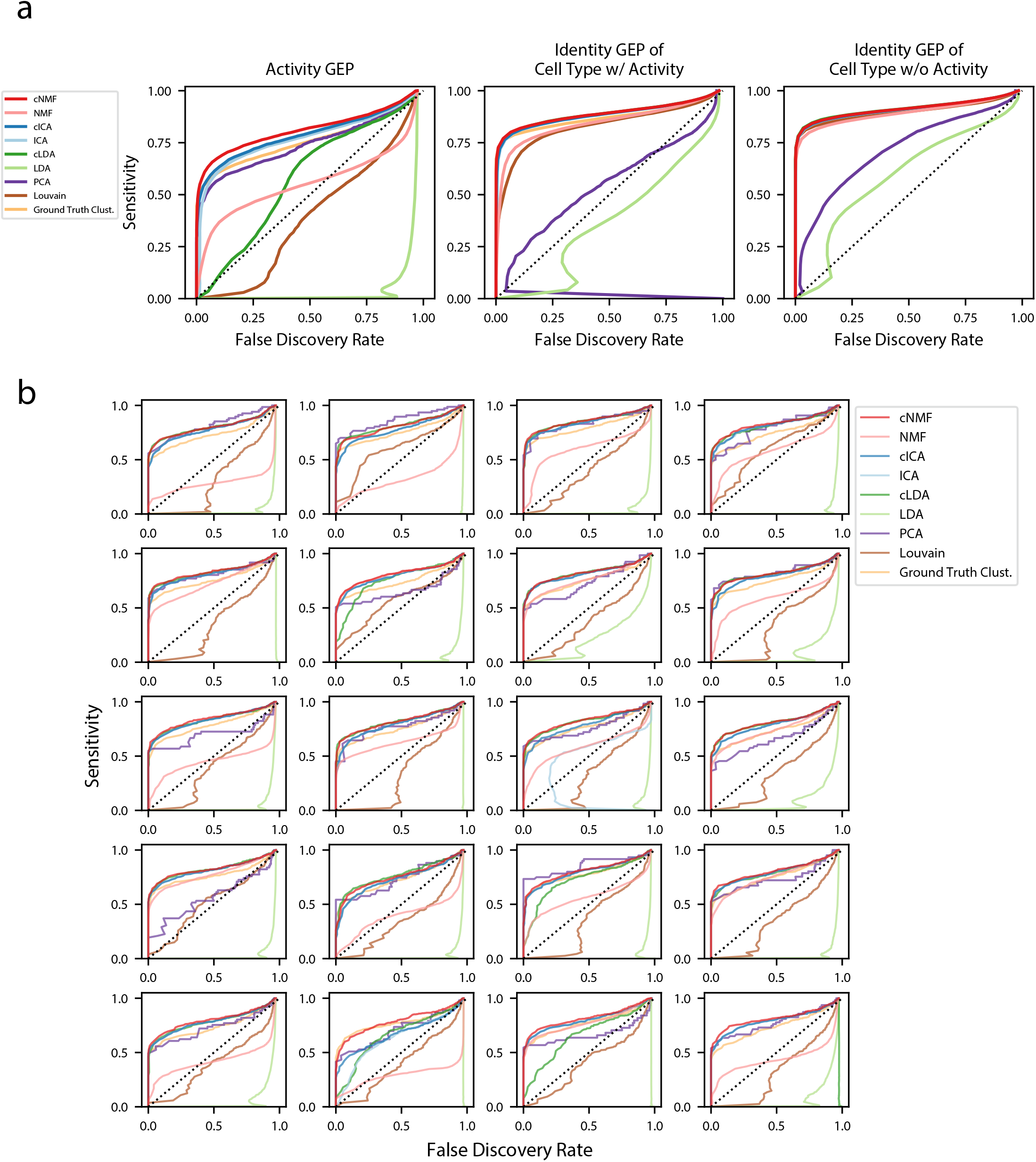
Accuracy of identifying genes in each GEP. **(a)** Receiver operating characteristics (ROCs) but showing false discovery rate (FDR) on the X-axis against sensitivity on the Y-axis for detection of genes in GEPs. Separate curves are shown for cNMF, cICA, cLDA, NMF, ICA, LDA, ground truth cluster assignment, Louvain clustering, or PCA. These show combined results for all 20 simulations with mean log2 fold-change = 1.00. Sensitivity is calculated considering genes with a differential expression fold-change of >=2 and FDR is calculated considering genes with no differential expression (fold change = 1) **(b)** Same as (a) but only showing the results for the activity GEP and plotted separately for each of the 20 simulations at the mean differential expression log2 fold-change of 1.00.

**Supplementary Figure 6:**
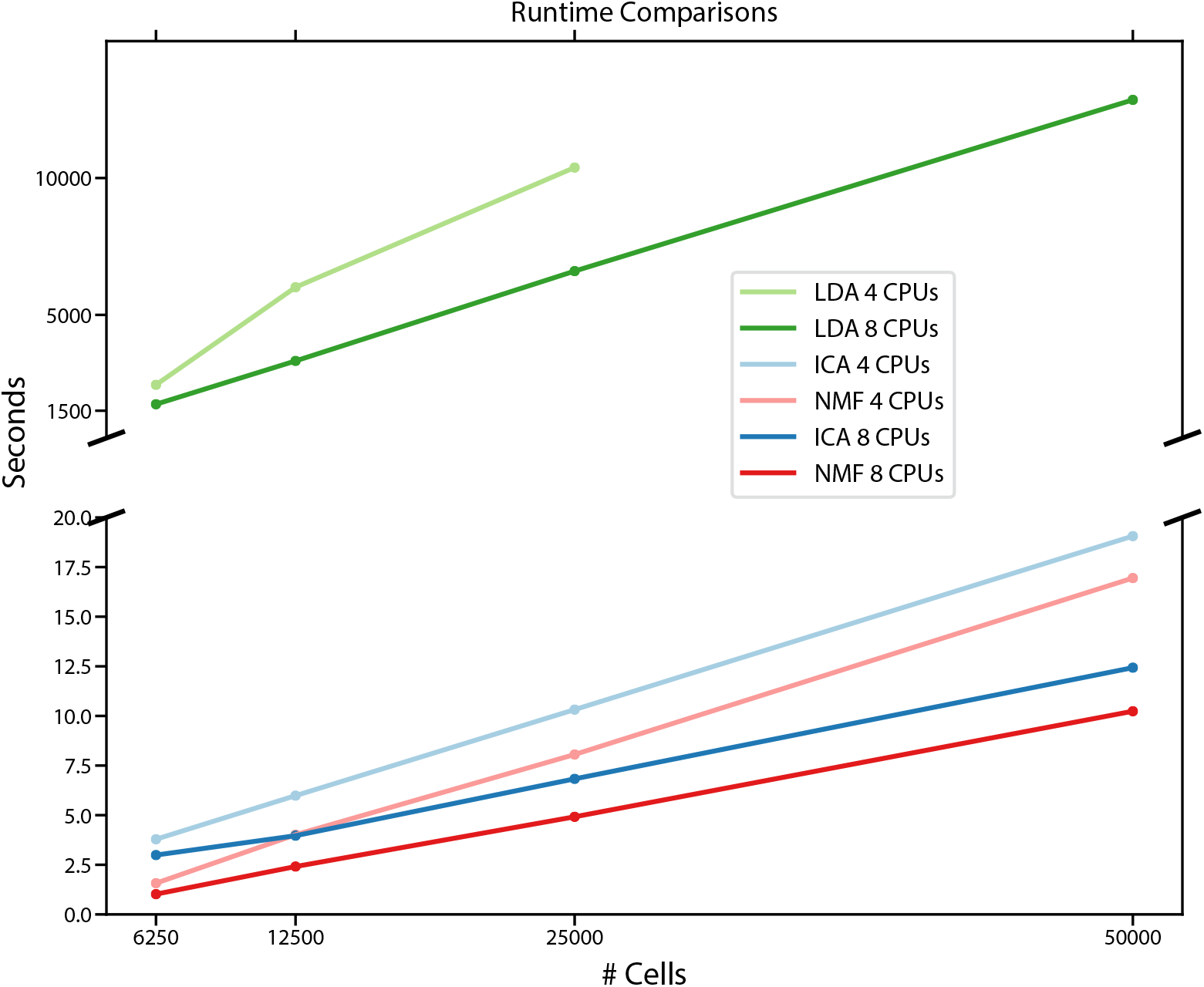
Comparison of run-times for different matrix factorization algorithms. Run-time in seconds for NMF, ICA, and LDA for a simulated scRNA-Seq dataset down-sampled to 6250, 12500, 25000, or 50000 cells, run either using 8 CPUs or 4 CPUs. Estimates are the average of 3 independent replicates with different seeds.

**Supplementary Figure 7:**
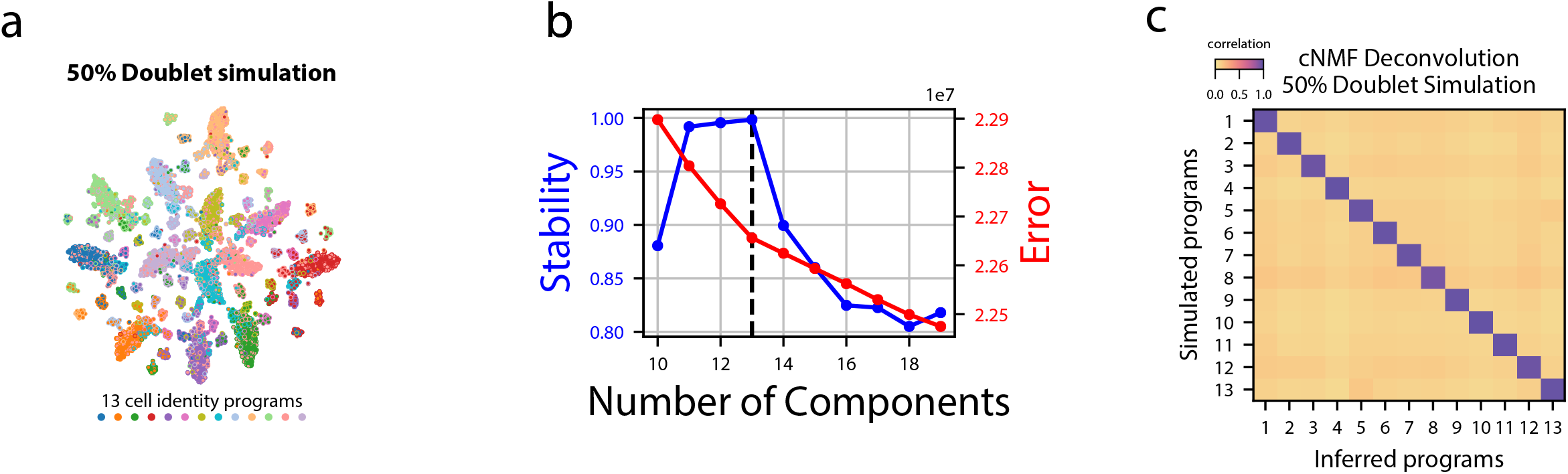
cNMF demonstration on simulated datasets with many doublets. **(a)** tSNE plot for a simulated dataset containing 50% doublets with marker color and edge color representing the simulated cell types. **(b)** K selection diagnostic plot showing solution stability (measured by the silhouette distance) in blue and Frobenius error of the consensus solution in red. **(c)** Pairwise Pearson correlation between ground truth GEP means (rows) and GEPs inferred by cNMF (columns) for the 50% doublet simulated dataset.

**Supplementary Figure 8:**
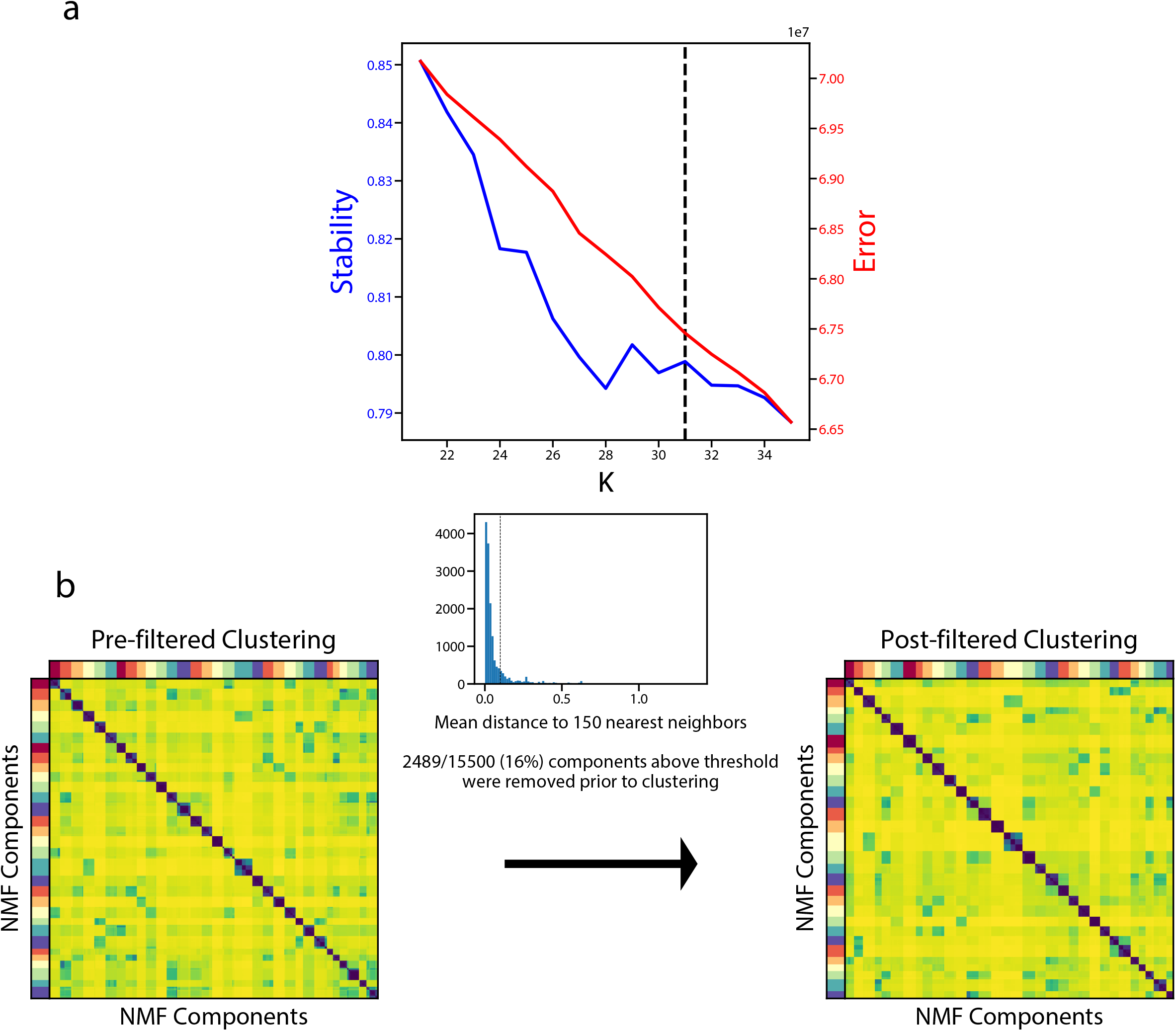
Diagnostic plot for cNMF on the Quadrato et. al., 2017 brain organoid dataset. **(a)** Number of cNMF components (K) against solution stability (blue, left axis) measured by the silhouette score of the clustering, and Frobenius error of consensus solution (red, right axis). **(b)** Clustergram showing the clustered NMF components for K=31, combined across 500 replicates, before (left) and after filtering (right). In between, we show the average distance of each component to its 150 nearest neighbors as a histogram with a dashed line where we set the threshold for filtering outliers.

**Supplementary Figure 9:**
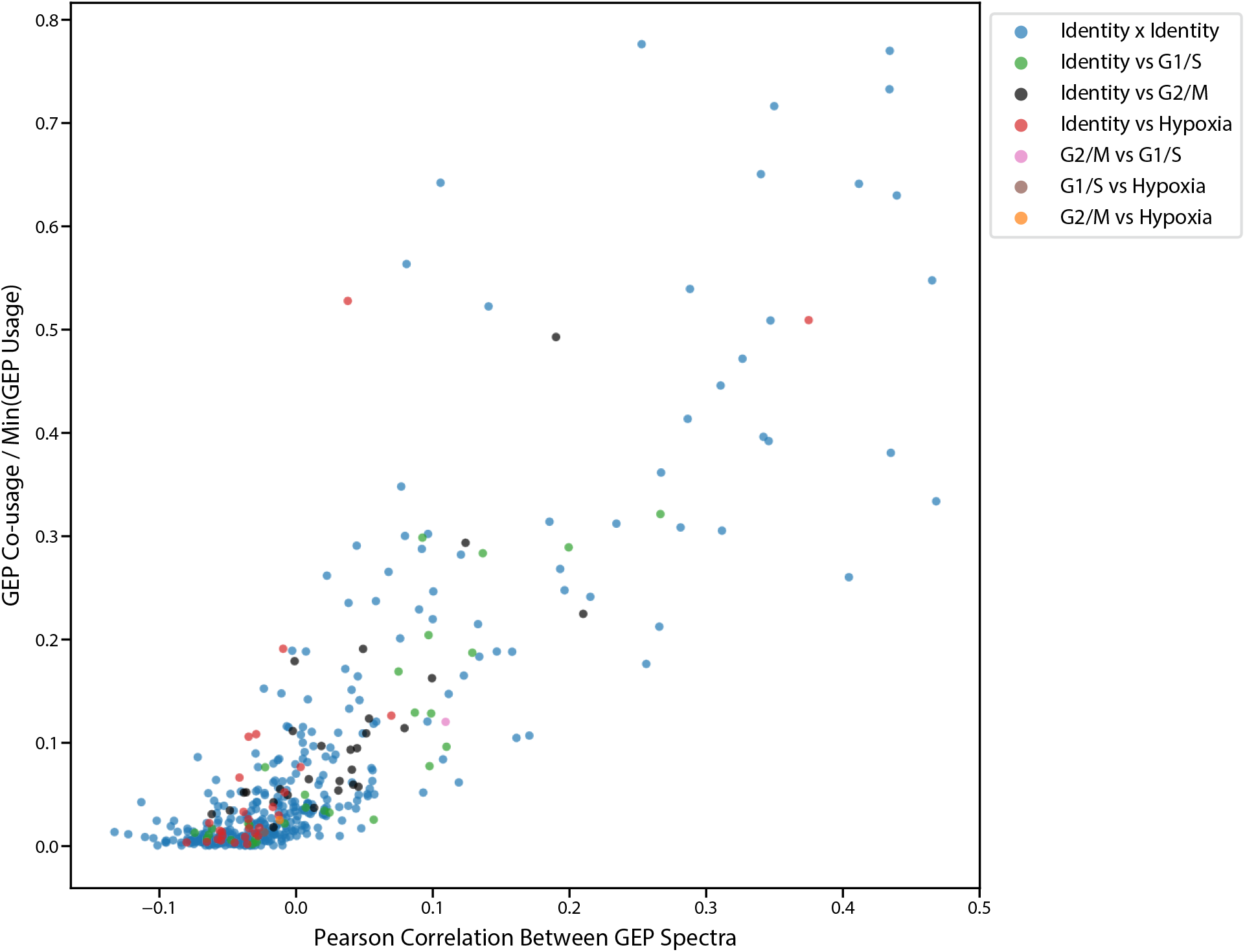
Correlation between GEP spectra pairs and fraction of cells that use both programs. Scatter plot of the pearson correlation between pairs of GEPs (X axis) and the fraction of cells that co-use the GEP pair (Y axis). Co-usage is defined as the number of cells with usage > .1 for both programs divided by the number of cells that use the less common of the programs with usage >.1. Dots are colored by whether or not the GEP pair is made up of identity or any of the 3 activity programs.

**Supplementary Figure 10:**
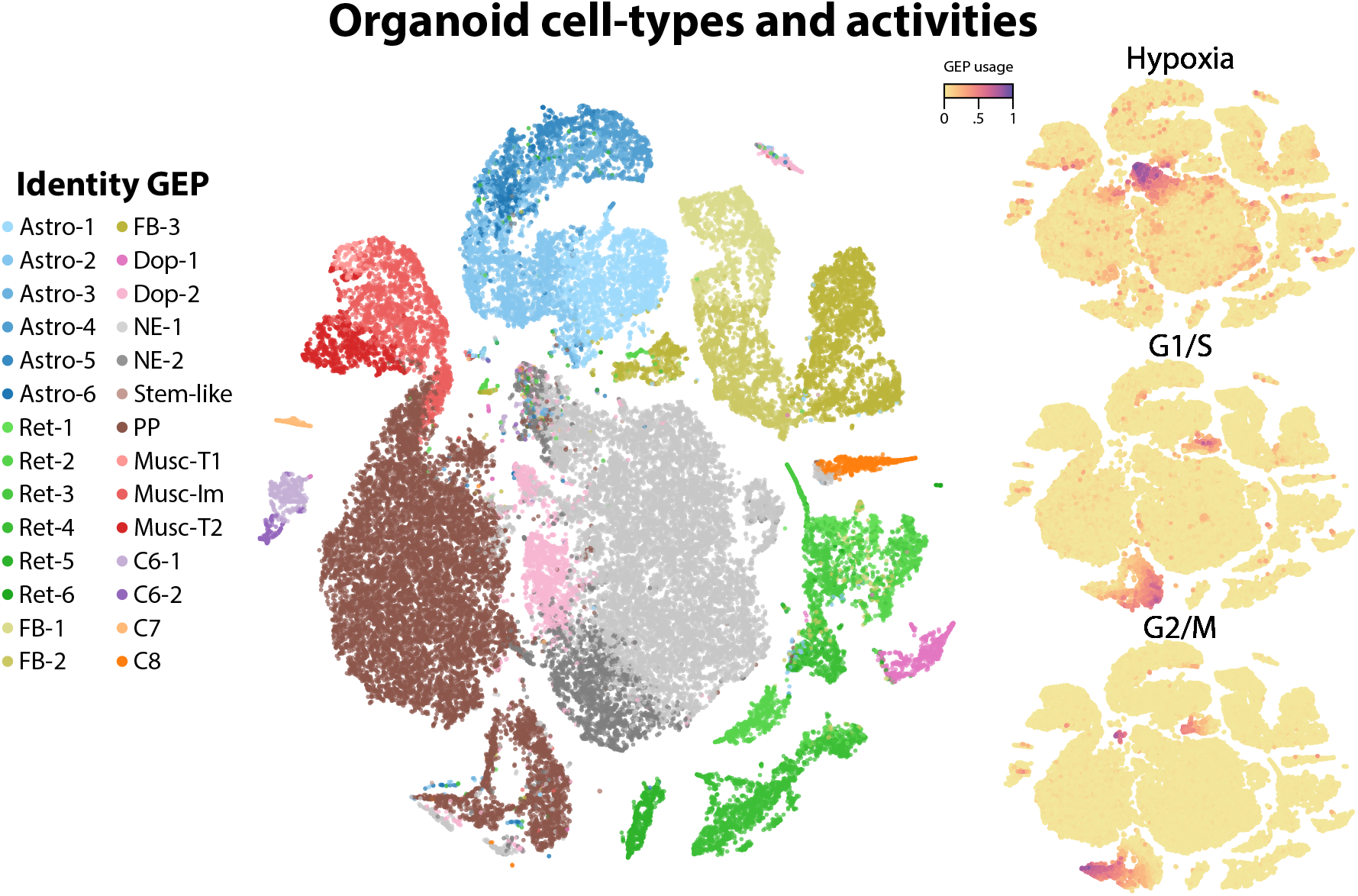
t-SNE plots of identity and activity GEPs in the Quadrato et. al. brain organoid dataset. t-SNE plots of cells colored by maximum identity GEP usage (left) or by absolute usage of each activity GEP (right).

**Supplementary Figure 11:**
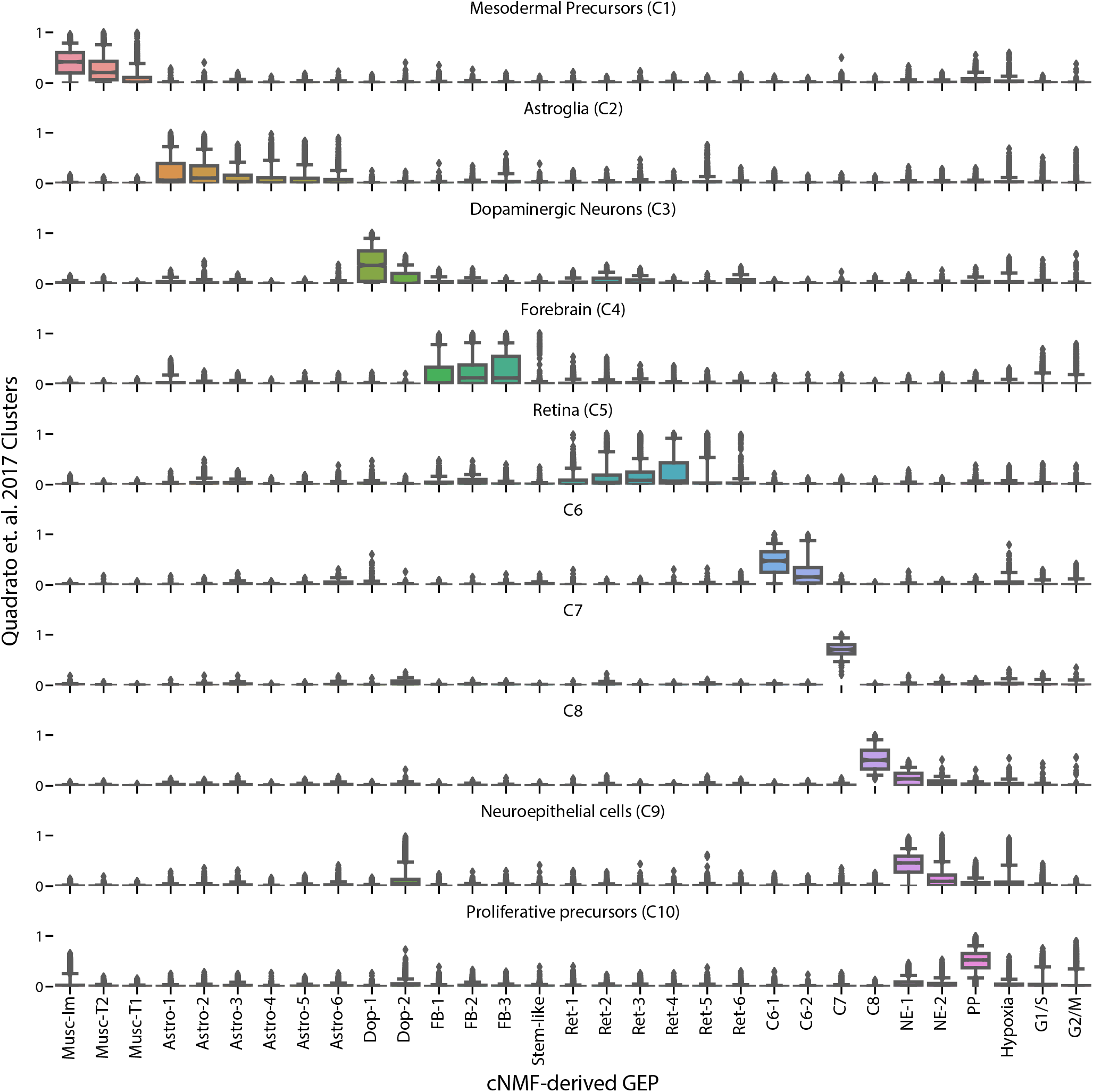
Comparison of cNMF usages with the cell-type clusters from Quadrato et. al., 2017. Box and whisker plot of the usage of each GEP (column) in cells of the clusters from Quadrato et. al. 2017 (rows). Boxes represent interquartile range, whiskers represent 5th and 95th percentiles.

**Supplementary Figure 12:**
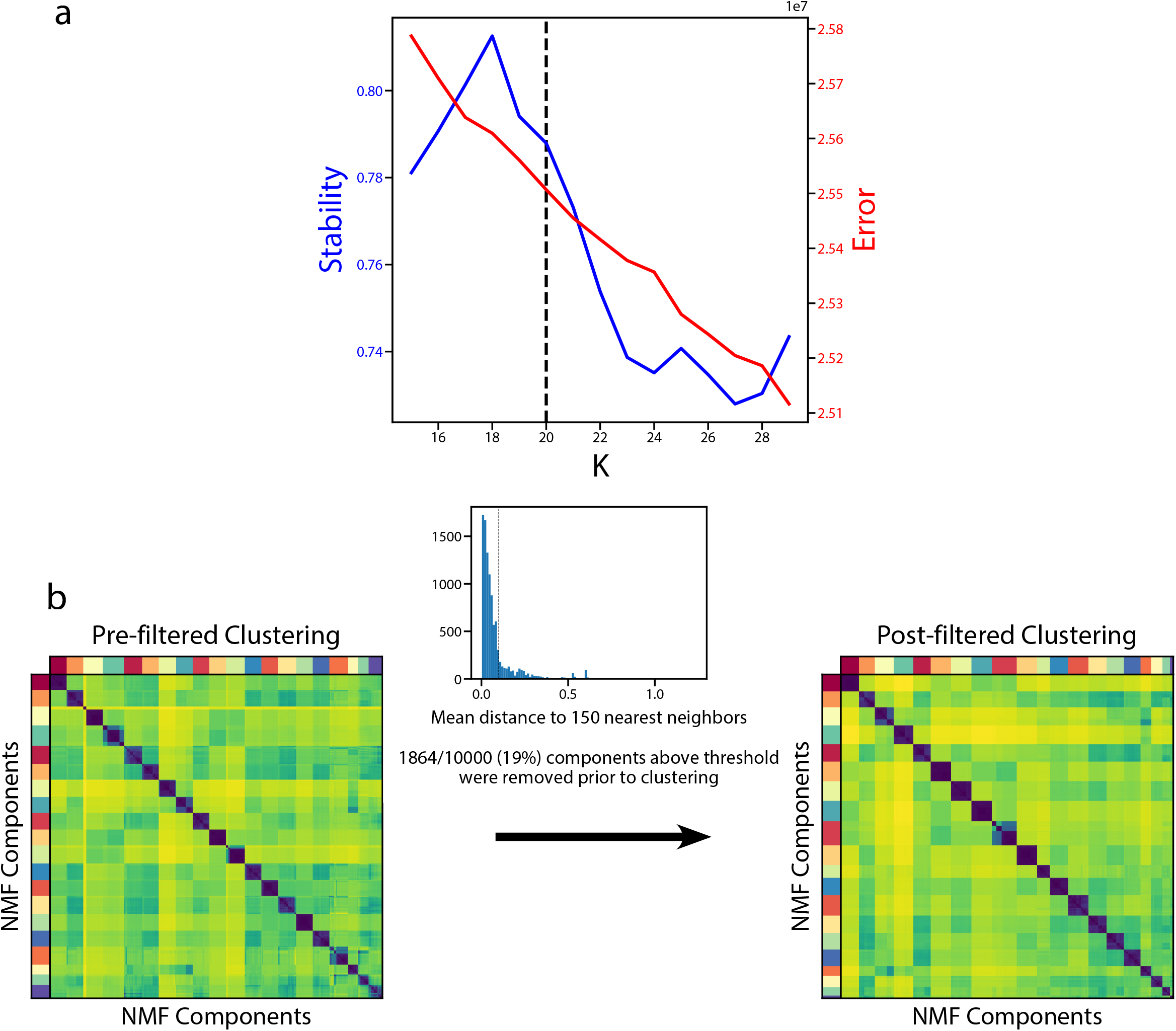
Diagnostic plot for cNMF on the Hrvatin et. al., 2017 visual cortex dataset. **(a)** Number of cNMF components (K) against solution stability (blue, left axis) measured by the silhouette score and Frobenius error of consensus solution (red, right axis). **(b)** Clustergram showing the clustered NMF components for K=20, combined across 500 replicates, before (left) and after filtering (right). In between, we show the average distance of each component to its 150 nearest neighbors as a histogram with a dashed line for the threshold for filtering outliers.

**Supplementary Figure 13:**
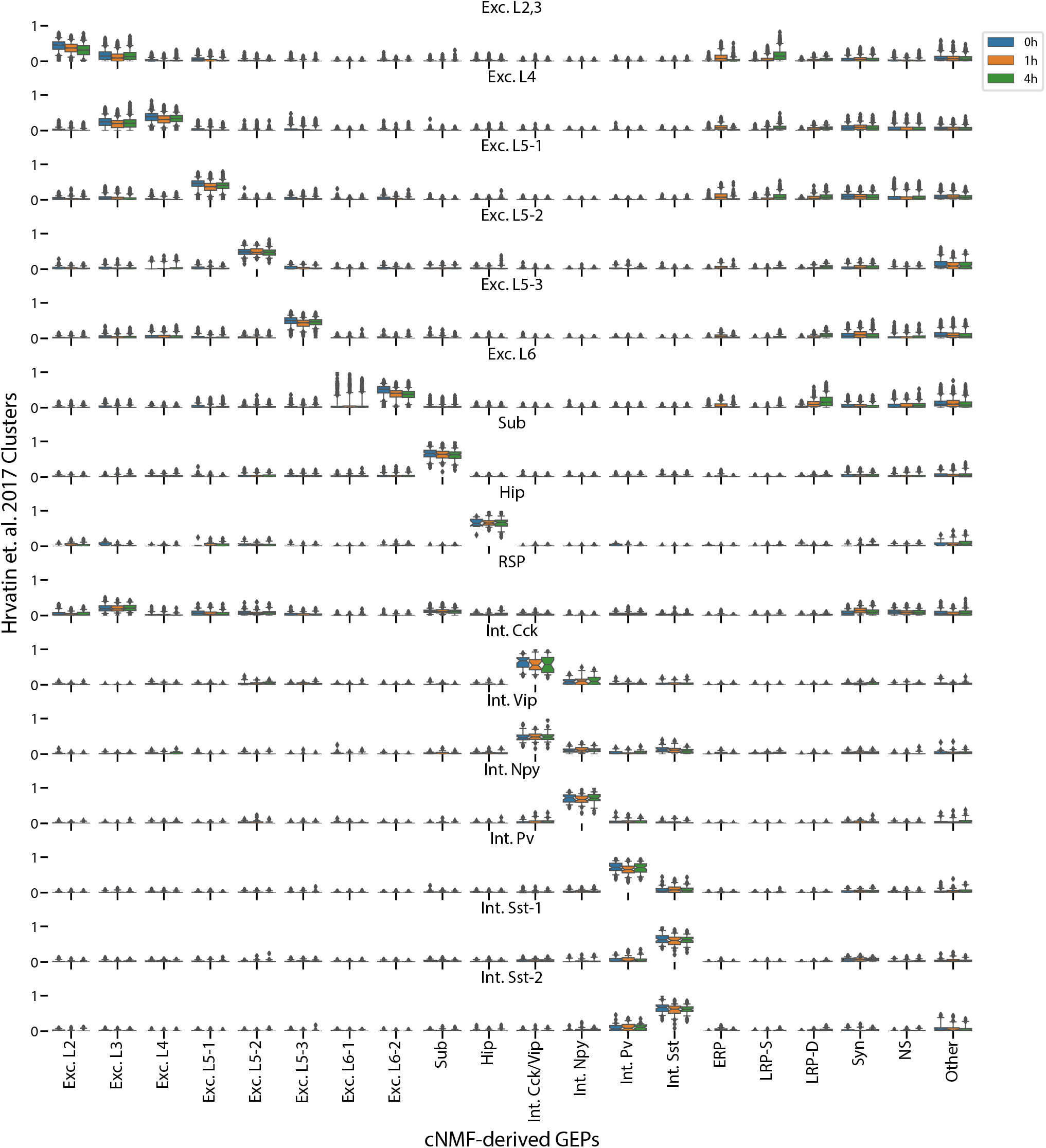
Comparison of cNMF with the visual cortex cell-type clusters from Hrvatin et. al., 2017. Box and whisker plot of the usage of each GEP (column) in cells of each cluster from Hrvatin et. al. 2017 (rows) stratified by the stimulus condition of those cells (hue). Boxes represent interquartile range, whiskers represent 5th and 95th percentiles.

**Supplementary Figure 14:**
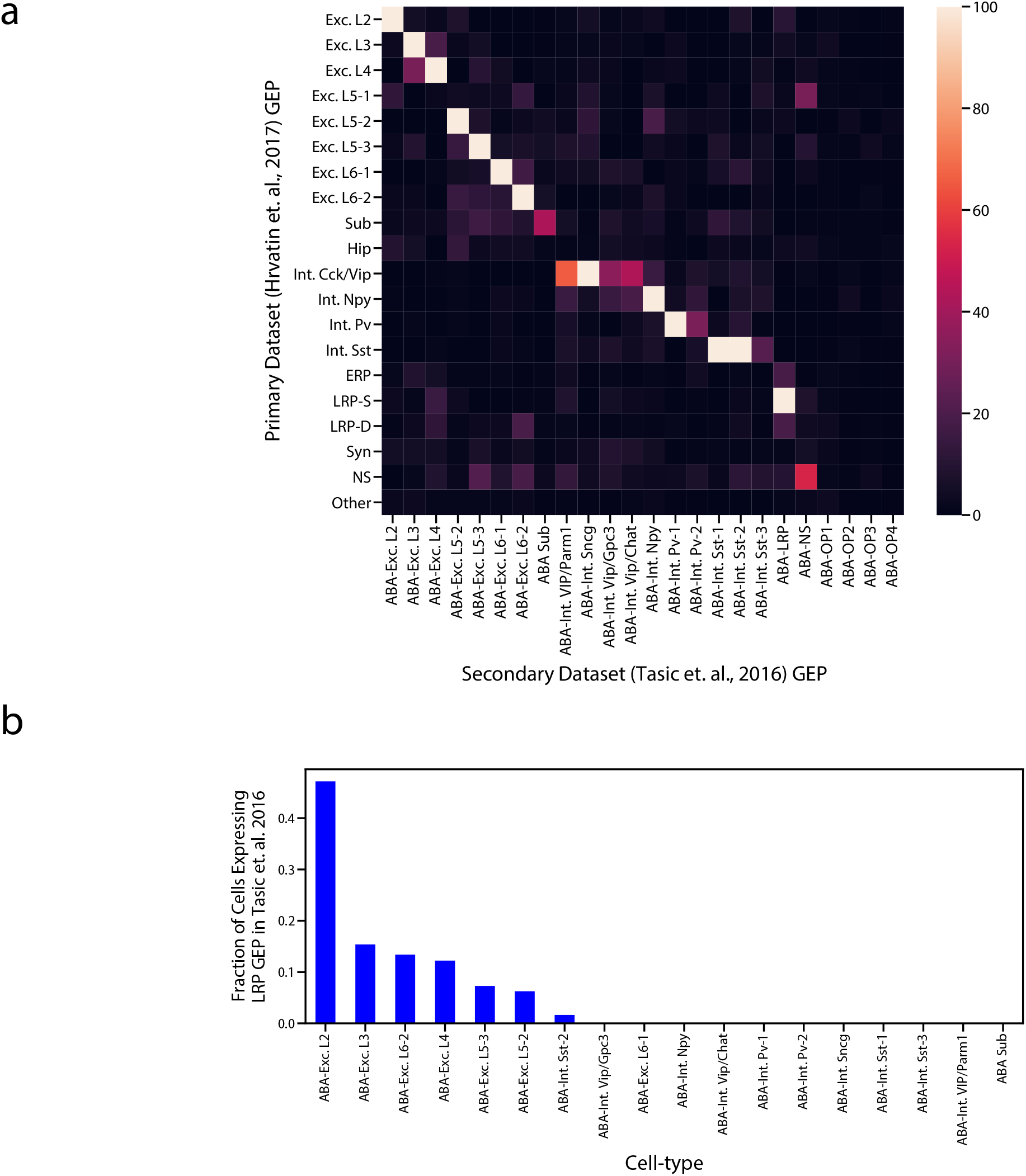
Comparison of GEPs identified in the Hrvatin et. al., 2017 and Tasic et. al., 2016 visual cortex datasets. **(a)** Heatmap showing the odds ratio for the intersection of top associated genes in each inferred GEP in the Hrvatin et. al., 2017 and Tasic et. al., 2016 datasets. Top associated genes were defined as those with an association score >= 0.0015. Odds ratios above 100 were set to 100 for better visualization of pairs in the lower range. GEPs from the Tasic et. al. dataset are labeled as ABA for Allen Brain Atlas. **(b)** Proportion of cells of each cell type that express the superficial LRP with greater than 10% usage in the Tasic et. al. dataset. Cells were assigned to a cell type based on their most used identity GEP.

